# PloidyPeaks: R package for automated flow cytometry cell cycle analysis and visualization in fungal microbes

**DOI:** 10.64898/2026.10.29.762876

**Authors:** Margot Henry, Caitlin Kostna, Leanne von Giese, Aleeza Gerstein

**Author notes:** Address correspondence to Aleeza Gerstein,.

## Abstract

Karyotypic variation in fungal populations from ecological, clinical, and industrial environments and is also a hallmark of many cancers. The evolutionary dynamics of karyotypic vriants (i.e., changes in the number of chromosomes, either as complete sets or single chromosomes) are the same as any *de novo* mutation. To quantify karyotypic subpopulations within evolving populations, an unbiased computational method is required to identify subpopulations and quantify their abundance. Flow cytometry is the standard method to measure the genome size from large populations of individual cells. Cell populations typically include cells in all stages of the cell cycle (G0/G1 before DNA replication, S phase during replication, and G2/M after DNA replication but before cell division). Existing mathematical models can be fit to genome-size distributions to determine the base ploidy of a population, but existing algorithms assume a single base ploidy. Mixed-karyotype populations therefore require manual gating to subdivide the flow cytometry data before analysis, which introduces bias, as it is not always clear where or when gates should be drawn. To overcome this, we developed an new method that identifies and quantifies karyotypic subpopulations from flow cytometry data. The open-source R package *ploidyPeaks* builds on existing Bioconductor tools to import and manipulate standard *.fcs* flow cytometry files. Using reference populations of known ploidy, we established thresholds to distinguish single-genome size populations and identify candidate mixed-karyotype samples. We then developed a peak-detection algorithm to identify G0/G1 and G2/M peaks, resolve karyotypic subpopulations within mixed populations, and apply nonlinear least-squares fitting of the Dean-Jett-Fox cell-cycle model to each subpopulation, providing a confidence measure for model fit. Compared with existing methods, our approach quantifies genome size in mixed-karyotype populations without manual intervention while providing an objective measure of model fit. We demonstrate the utility of this method on 240 replicate populations initiated from 20 *Candida albicans* genetic backgrounds evolving for 100 generations to the antifungal drug fluconazole.

## INTRODUCTION

Karyotypic variation in ploidy (the number of complete chromosome sets) and aneuploidy (an aberrant number of chromosomes) has been observed among isolates of microbial fungal species that span the kingdom [1, 2, 3, 4]. Unlike sexually reproducing eukaryotes, these species predominantly undergo asexual reproduction (i.e., mitosis rather than meiosis), which removes a major checkpoint for the maintenance of euploidy. When cytokinesis fails after replication, this doubles the number of chromosome sets in one step (’endoreduplication’), creating diploid cells from haploids or tetraploids from diploids. Chromosomes can also be gained or lost due to errors in chromosome segregation due to improper spindle pole body function or misattachment of the spindle microtubules to the chromosomes [5]. Karyotypic mutants have been observed in many contexts, including isolates acquired from ecological, clinical, and industrial environments ([3, 5], and references within). There are some clear cases where aneuploidy has been linked to an adaptive trait (most commonly drug resistance, [6, 7, 8, 9, 10, 11, 12, 13, 14, 15], though also morphological transitions [16, 4]). However, the true incidence of karyotypic variation is natural populations is hard to estimate, as sampling natural populations is often restricted to only a small number of isolates per site, and the methods used to acquire pure isolates may select against variant cells [17].

Furthermore, isolating the fitness effect of karyotypic variation is not trivial compared to genic mutations, as we lack the corresponding molecular tools to precisely recreate karyotypic variants in a given otherwise isogenic background. Thus, the ecological and evolutionary influence and importance of karyotypic variation in natural populations evolving under diverse environmental conditions remains somewhat obscure. Simply observing karyotypic variation does not necessarily mean it is beneficial, as some environments, such as azole drug, also seem to induce aneuploidy as well as select it [18].

Experimental evolution studies (also sometimes referred to as “adaptive laboratory evolution”, ALE) have emerged as a powerful way to test how different factors influence the phenotypic effect and eventual evolutionary fate of karyotypic variants. In these studies, the researcher typically sets up a series of replicate populations from the same ancestor that are then evolved in the lab to a specific environment [19]. This permits direct studies on how different factors influence the rate of karyotypic variation formation and theoretically their corresponding fitness effects. The frequency of observed karyotypic variant cells in these experiments depends on the ancestral ploidy and the environment ([20, 21, 22]). In all cases, as with any *de novo* mutation, there is a period of time after the new karyotypic mutation appears in the population when the population is heterogeneous for karyotype, i.e., when multiple different subpopulations are observed. In order to understand the dynamics of karyotype subpopulations and their role in adapting populations, we require a method that can identify different subpopulations and quantify the number of cells within them.

Flow cytometry is the standard method to measure genome size (as a proxy for karyotype) from populations of cells. The cells are fixed and then stained with a fluorescent dye that binds to DNA. The flow cytometer is used to measure genome size from typically *→*10 000-50 000 cells in a population by passing them one-by-one across a laser. Typically cells are present in all phases of the cell cycle: G0/G1 (which we will refer to as G1) prior to DNA replication, S phase during replication, and G2/M (which we will refer to as G2) when cells have double the DNA but haven’t yet divided. The flow cytometer scatters light though each suspended particle, the light scatters are read into a detector that can visualize the light, and this information is passed on to an analyzer that converts them to electrical signals. Multiple parameters can be measured for each cell.

Physical characteristics such as cell size and shape are measured based on light scatter. Forward scatter (FCS) measures the light scatter in the same direction as the laser, providing information on cell size, while side scatter (SSC) measures scatter on a 90*^↑^* angle, providing information on cell granularity ([23]). When the cells get passed through the laser, it pulses in specific fluorescence channels that are named based on machine settings, and the height (-H), area (-A), and width (-W) are recorded. The fluorescence intensity parameters are typically labelled by arbitrary fluorescence/laser numbers (e.g., FL1, FL-2, etc.) or the name of the dye used to stain the DNA (e.g., FITC, PI, SYBR, SYTOX etc.). Genome size (a composite measure of ploidy and aneuploidy) correlates with the fluorescence intensity of the G1 phase cells. Genome size from cell populations of interest are compared to the G1 mean from control populations where the ploidy is known.

Software such as Flow-Jo (Treestar) or Floreada.io fit a cell cycle algorithm to the fluorescence intensity data to determine G1 mean. There are two different cell cycle algorithms that are built-in to Flow-Jo to measure genome size, Watson Pragmatic ([24]) and Dean-Jett-Fox [25]. They have generally the same process; they both assume that cells in the G1 and G2 clusters are Gaussian distributed but vary in how they fit the distribution of S phase cells. With fungal microbial populations, the two models often give identical or near-identical results. Model fit can be evaluated by visual comparison by comparing curves of the fitted model overlay on top of a histogram of fluorescence. The cell cycle algorithms are defined to identify peaks from a single ploidy population and either fail when there are subpopulations (poor fit between model and fluorescence histogram) or ignore prominent peaks in the distribution. One work around has been to manually draw gates to divide the original population by eye into subpopulations and then run the cell cycle algorithm on each. Although this seems to work to some degree some of the time, it is very time-consuming, is subject to bias (as you will only find the populations that you have identified by eye, relying on intuition when looking at the raw data), and is not feasible when there are many subpopulations, or without prior experience with flow data.

Other analysis packages exist to quantify ploidy from flow cytometry data through other algorithms. *flowClust* uses a model-based clustering approach to determine the number of clusters within the sample. The package estimates the number of clusters by Bayesian Information Criterion and the Integrated Completed Likelihood ([26]) to provide the total number of clusters within the sample. The package does not provide information about G1/G2 subpopulation pairings explicitly, making it difficult to infer subpopulation information. The package *flowPloidy* uses a non-linear regression model of DNA fluorescence intensity to find the cell populations that contribute to the total sample data to determine ploidy ([27]). The G1 and G2 peaks are modelled as normal distributions and the regression model estimates the means of each. The function FlowHist() creates a histogram and applies the model. FlowHist() requires some user input on population information. The function assumes there are two subpopulations; if there are more or fewer the user must indicate otherwise. There is a logical parameter called *g2*, which defaults is true, used to indicate if each sample has a G2 peak. There are some samples that will only have one peak (i.e., no corresponding G2 peak), which requires the user to change the parameter flag. The model is sensitive to local minima traps, when the non-linear regression model misclassifies a local minima as the G1 or G2 peak ([27]). The solution was the creation of a function called browseFlowHist(), a graphical user interface (GUI) that allows the user to correct and review the model applied to the histograms manually. If the model fails or does not correctly identify the peaks, this function allows the users to use their mouse and click on the GUI to properly identify the peaks.

The current software does not consider the potential presence of doublets, which can complicate the assessment of subpopulations. Doublets are an excess of cells that happen when two cells pass through the laser at the same time and can be mistaken for a cell population (G1 or G2) of high genome size ([28]). Doublets typically appear around the G1+G2 range and sometimes around the G2+G2 range. In the case of a mixed population that contains a subpopulation created through endoduplication, the new population will have a G1 phase peak equal to the G2 peak of the original population, with the second G2 peak at G2+G2 of the original population. Thus it is also not trivial to always discern which peaks correspond to doublets (and can be effectively ignored) and which peaks are indicative of a subpopulation.

Our objective was thus to develop an open-source method to quantify karyotypic variation in populations from DNA fluorescence intensity data in an unbiased manner that removes user input as much as possible. We hence sought a method that would be able to find all of the peaks in a sample, is able to identify peaks corresponding to potential subpopulations (i.e., G1/G2 pairs), and to make inferences about the presence of doublets. In addition, we wanted to provide a measure of model fit for each population to create a confidence term to provide the user a sense of how well the algorithm performed. Since visualization is an important component of the flow cytometry workflow and analysis, we also wanted to create methods to visualize a large number of samples simultaneously since many of our experiments are done with hundreds of samples. To accomplish this, we developed an R package, *ploidyPeaks*, that uses peak detection algorithms based on the histograms of DNA fluorescence intensity data to identify potential G1 and G2 peaks. We consider every local maxima in the fluorescence intensity data to be a possible peak, and then use the Dean-Jett-Fox cell cycle algorithm to identify G1/G2 subpopulation pairings without requiring user input. A confidence term is calculated using the nonlinear least squares method to calculate a residual standard error, indicating how well the final model fits the data.

## MATERIALS AND METHODS

### Nomenclature

We follow the standard flow cytometry nomenclature. A sample is the data from a single population of cells. The information on all variables meaured (e.g., FCS, SSC, FITC) is contained in a single *.fcs* file. Samples can contain cells with one karyotype (i.e., containing single G1 and G2 peaks) or multiple subpopulations (multiple G1/G2 peaks). The phrasing ‘flow frame’, used by some packages, is synonymous with a single sample. A flow set is a group for samples (or flow frames), composed of multiple *.fcs* files. The data from a flow set is typically collected on the same day and stored together in a single folder. A flow cytometry sample contains data from single cells, and often also contains spurious measurements from small particles (debris) with small size and low fluorescence, and sometimes large clusters of cells with high size and high fluorescence. The first step of a flow cytometry analysis workflow is to apply what is termed “a gate” to retain only the cells inside the boundary for further analysis.

### Import and gate data

*ploidyPeaks* builds off the *flowCore* Bioconductor package; *flowCore* has over 100 functions to bring in, manipulate, transform, and save flow cytometry data in the R programming language. The functions *read.FCS()* and *read.flowSet()* are used to import data from a flow frame or a flow set, respectively, to ensure the provided data are in the correct *.fcs* file format, and to save the information into the S4 structure that is required for Bioconductor packages.

Following standard flow cytometry practices, gating in *ploidyPeaks* creates boundaries on a bivariate density plot with SSC-A on the y-axis and the fluorescent dye intensity on the x-axis. This creates a rectangle that will retain the cells within the boundaries and remove all other cells from the subsequent analyses. The goal is to include the majority of cells, excluding only those truly on the margins. The gating functions *rectGateFlowFrame()* and *rectGateFlowSet()* were written to apply gates to either a flow frame or flow set. Both functions combine existing *flowCore* functions: *rectangleGate()* is used to specify the gate, *Subset()* retains the cells within, and *write.FCS()* saves the gated data for further analysis. We also employ *autoplot()* from the *ggcyto* package to visualize the bivariate density plot, *geom_gate()* to add a red rectangle on the plot to visualize the gate, and *as.ggplot()* to visualize plots side by side (Figure **1**).

**FIG 1.**
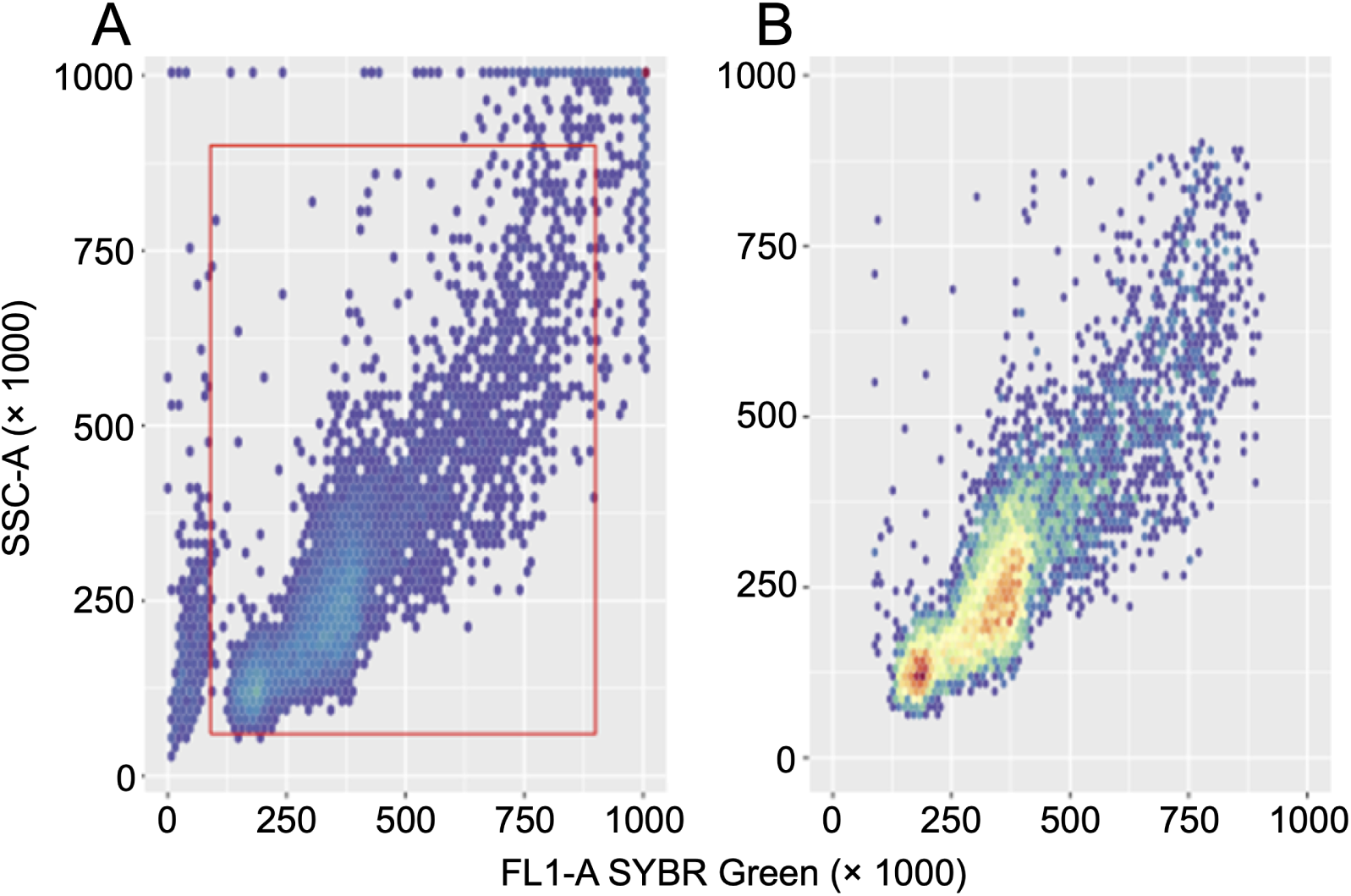
Gating using a bivariate density plot. The x-axis is the fluorescence intensity data (in this case Sybr Green was used, which was read in the FL1-A channel), on the y-axis is side scatter area (SSC-A). Each point on the bivariate plot represents a single cell, density is indicated by a blue (low) to red (high) cell density gradient. A) The red rectangle is the gate applied to the sample. B) Cells within the gate are retained and used for further analysis. Note that the magnitude of numbers is arbitrary and determined by machine settings.

### Identify and G1 and G2 peaks in single karyotype population samples

The core goal of *ploidyPeaks* is to determine whether samples contain a single karyotype population or multiple subpopulations and to identify all corresponding G1 and G2 peaks. This is done through analysis of the fluorescence intensity data, corresponding to genome size, from each sample. This data can be visualized as a histogram of the underlying cell-level data, with the peaks indicating high cell count values. The distribution of cells varies depending on the underlying population (Figure **2**).

**FIG 2.**
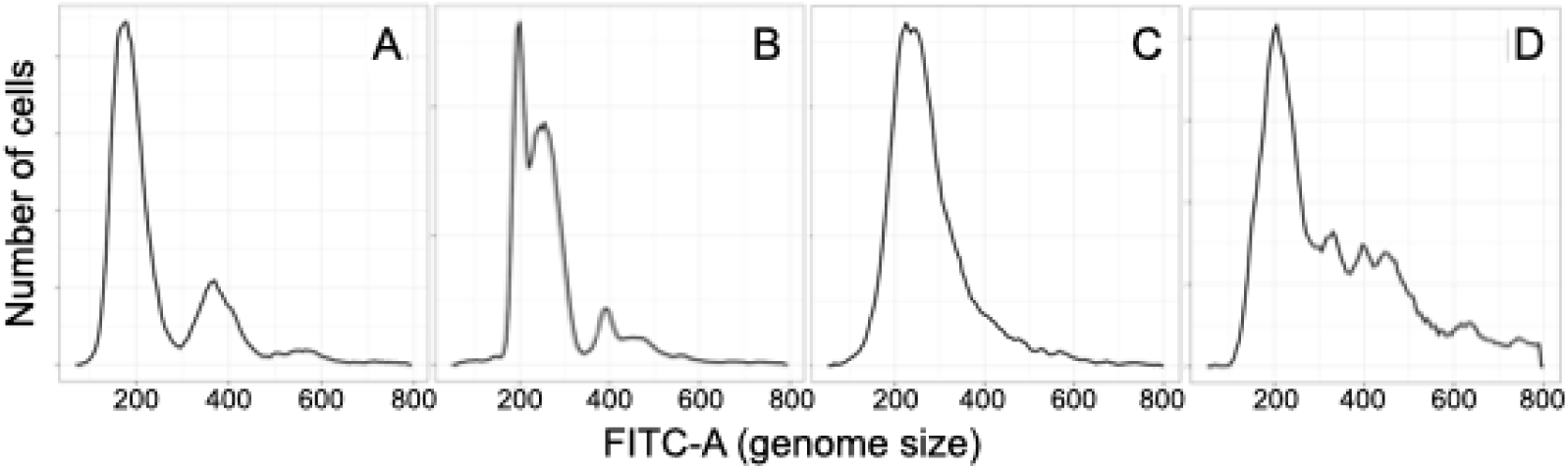
Examples of different shapes and sizes of genome size peaks. A) Some samples have two obvious G1 and G2 peaks. B) Some samples have multiple distinct peaks. C) Some samples have only a single distinguishable peak. D) Some samples have atypical distributions where the association between G1 and G2 peaks is not obvious.

A single wrapper function called *flowPeakDetection()* analyzes the data and categorizes each sample as a single population or multiple subpopulations. *flowPeakDetection()* iterates through a series of algorithms, *peakAlgorithm1()* – *peakAlgorithm5()*, to identify G1 and G2 peaks and pair them when appropriate (Figure **??**). The first step in all algorithms is identifying local maximums (peaks, as visualized on a histogram) in the univariate fluorescence intensity data. There are typically small spikes in the distribution of DNA fluorescence values caused by stochasticity in the cell population; if all local maxima are retained this would yield many peaks that do not reflect the true biological variation.

These excess small peaks are removed by using the *rollmean()* function from the *zoo* package [29] to smooth the data. The moving mean, also called the rolling mean, calculates a series of means over a specified interval. This effectively eliminates the stochastic small spikes since neighbouring values average them out, while true peaks are retained (Figure **??**A & B). *peakAlgorithm1* uses an interval (’smoothing level’) of 11, which is effective for samples that contain a single population. Once the data is smoothed, the function *detect_localmaxima()* from the *scorepeak* package is used to detect local maxima in a given width for univariate data [30]. The *scorepeak* package was developed to identify true peaks from false peaks created by noise in times series data [30]. We use the reflecting boundary condition, which considers the neighbouring values to the right and to the left of each value of *x_i_* in a given sliding window size (n) to decide if there is a peak. The smallest possible value for a sliding window (*n* = 3) is specified to ensure that each potential genome size peak will be identified. The function *findTruePeaks()* is then called by *peakAlgorithm1*, as another layer to further remove spurious peaks caused by stochasticity. Testing datasets were used to find the average distance between peaks identified in *Candida albicans* samples with known subpopulations that had similar genome sizes, i.e., multiple G1 peaks close together, to determine the minimum distance where we can confidently say that multiple true peaks are present. If two clusters are too close together (i.e., below that distance), they are combined, and only one peak is kept per cluster.

If more than two peaks remain after *findTruePeaks()*, the peak that is furthest left (smallest genome size) is marked as a true G1 peak, and *flowPeakDetection()* searches for a matching G2 peak within specified boundary conditions. The boundary conditions build off the Watson Pragmatic (WP, [24]) and the Dean-Jett-Fox (DJF, [25]) cell cycle algorithms. They both identify the tallest fluorescence intensity peak and assign it the G1 mean. WP identifies the location of the highest point near 1.75*×*G1 mean to find the G2 mean, while DJF identifies the location of the highest point near the 2*×*G1 mean to find the G2 mean. We set the boundary conditions for *peakAlgorithm1* at (1.75 *× G*1, 2.2 *× G*2). Having identified G1 and G2 peaks, the population boundary is calculated at the location of the G2 peak plus half the distance between the G1 and G2 peaks (Figure **3**). The percent of cells outside the boundary (*popBoundary*) is calculated. If it is below the specified threshold (which defaults to 8.5%, determined through the testing dataset), the population is marked as a single population. If the *popBoundary* is greater than the threshold, *peakAlgorithm1()* fails.

**FIG 3.**
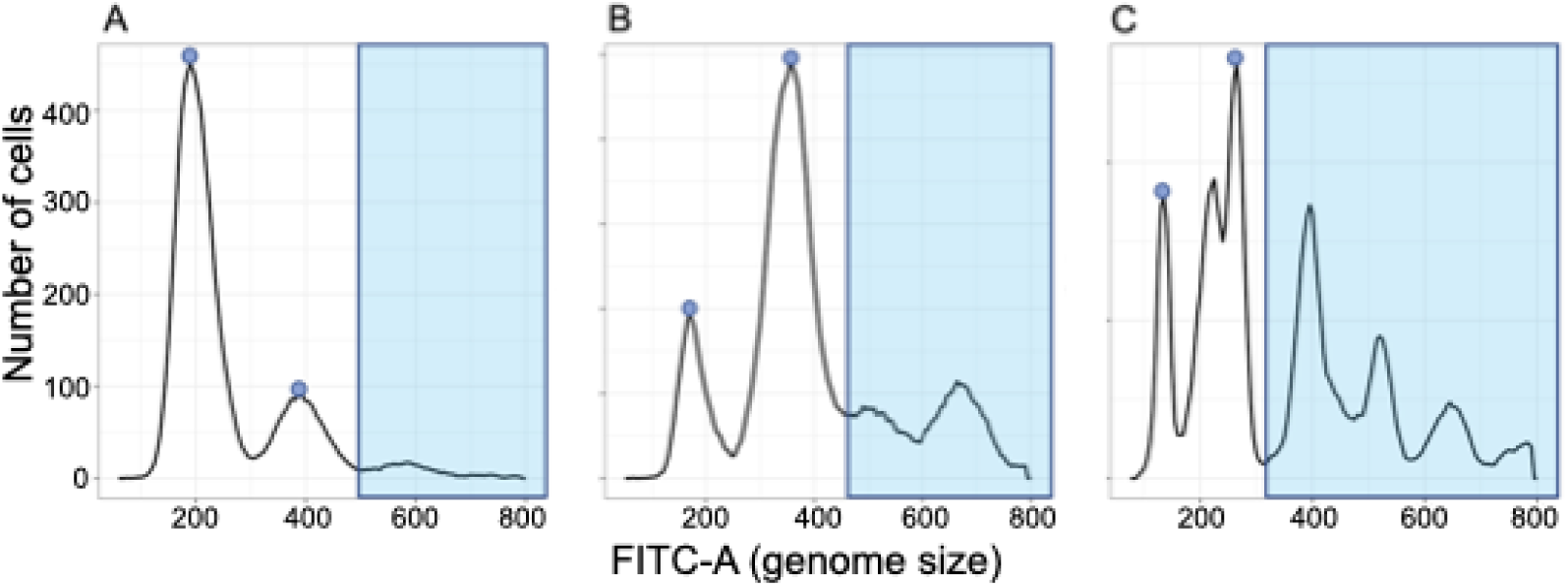
Example of the *popBoundary* on different samples. *PeakAlgorithm1()* calculates the percentage of cells to the right of the G2 peak (i.e., beyond *popBoundary*). The blue dots represent the G1 and G2 peaks used to find *popBoundary* and the blue shaded area represents the cells found in *popBoundary* A) An example of data from a single population sample, with only 5.5% of cells beyond *popBoundary*. B) An example of a population that would be passed on to *PeakAlgorithm2()*, where the percent of cells (32.4%) exceeds the set 8.5% threshold. The peaks in the boundary region are not called as doublets since the peak height at G2+G2 is higher than the peak at G1+G2 C) An example of a population with multiple populations. The algorithm assigns a G1 and G2 pairing from the peaks that are farthest to the left as the focal population. This sample would also be passed on to *PeakAlgorithm2()*, as the percent of cells in *popBoundary* (49.10%) exceeds the set threshold.

Occasionally the peaks identified using the peak detection algorithm weren’t always at the left most spot-specifically with the G1 peaks. Margot had implemented a flagging system for samples where the proportion of cells to the right of the G2 peak was excessive. This flag is of benefit to the user so they can go back and analyze the flagged samples individually. We decided that implementing a similar flag would be the best solution to our misidentified peaks. After some testing with some samples at various time points, Margot determined an appropriate threshold. The additional flag is used if the proportion of cells to the left of the G1 peak is below this specified threshold. A low proportion indicates that the G1 peak identified is most likely the actual G2 peak.

Samples can fail *peakAlgorithm1* for two non-exclusive reasons. The sample could contain multiple subpopulations, and the single sample model misses peaks representing cells in addition subpopulations. Single or multiple subpopulation samples may also contain additional peaks composed of doublets. Since the doublet peak(s) do not belong to any subpopulation, doublets increase the number of cells outside the population boundary, and when there are large doublet populations the peak algorithm will fail (Figure **4**).

**FIG 4.**
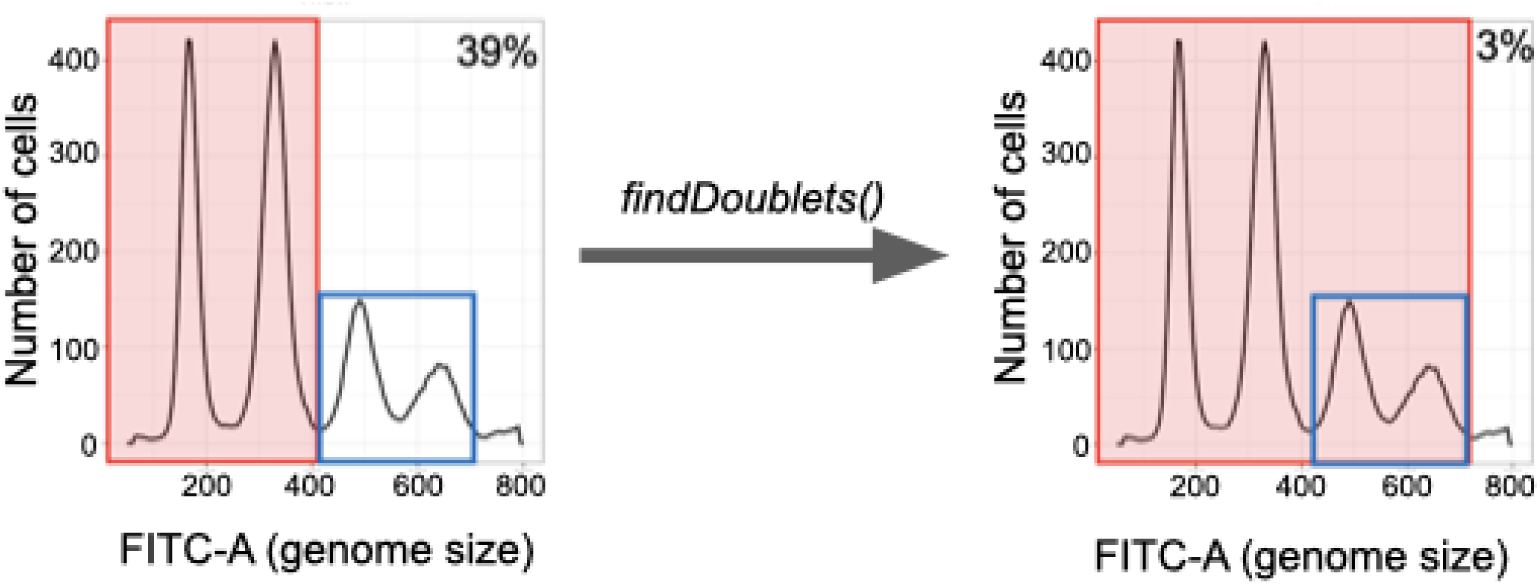
Identification of a G1+G2 anda G2+G2doublet in a single population sample. )T the sample would initially fail peak Algorithm1,since the proportion of cells being used outside the population boundary(cells in the whitespace)is 39%.B)textitfinddoublets()identifies the two peaks outlined in blue as G1+G2 and G2+G2 doublet peaks,which then accounts for 97% of cells and the sample passes.

The function *doubletCheck()* identifies potential doublet peaks in each sample. It looks for peaks around the G1+G2 range and the G2+G2 range. If there is a peak identified in that range, information is retained on the location and height of the doublet. There is unfortunately no known computational way to discriminate between a G1+G1 doublet and a true G2 peak. Similarly, the G2+G2 doublet can be mistaken as a G2 of an additional subpopulation. This becomes problematic in the case of populations containing an endoduplication subpopulation, where e.g., the 2N G2 peak is the same as the 4N G1 peak. Based on examination of a large number of flow samples single and mixed populations of known ploidy, we identified criteria to discriminate between doublets and a subpopulation G2. When doublets are present, the G1+G2 peak is always larger than the G2+G2 peak. This is consistent with a general observation that the G2 peak is rarely higher than the G1 peak. The G1+G2 peak is easier to classify as a doublet peak since there is rarely a corresponding appropriate G1 peak. Hence, if a sample contains a peak that could be assigned as either a subpopulation G2 peak or a G2+G2 peak, if that peak is smaller than a peak identified at the G1+G2 position, it is classified as a doublet(Figure **4** B). We also calculate an additional parameter, termed *maxDoubletHeight*, which calculates the height changes from G1 to G2 peaks. If the user notices their data has higher doublets or lower G2 peaks, *maxDoubletHeight* can be adjusted accordingly. Although we generally sought to reduce user input, nothing can make up for close examination of the specific datasets someone routinely collects. Data on doublets is retained analysis spreadsheet that is provided following analysis. To our knowledge, the existence of doublets within samples has previously been ignored, yet they may reflect something of biological interest since they are likely related to cell surface properties such as ‘sticky’ proteins that cause cells to stay together even through mechanical disruption. If doublets are identified within the sample, these peaks are accounted for, and *popBoundary* is calculated again. If the sample is again identified as likely containing multiple populations, and the data is passed to the next algorithm, *peakAlgorithm2()*.

### Identify and match G1 and G2 peaks in samples with subpopulations

The additional algorithms in *!*flowPeakDetection() re-initiate analysis at the peak detection step. In *PeakAlgorithm2()*, the smoothing factor is reduced to 5, which retains more granular data (i.e., more peaks and maintained), and the local maxima are again found. An additional step is added, to identify G1 and G2 pairs using the functions *findTruePeaks()* and an additional function, *findPairs()*. If the total number of identified peaks is even, this indicates that each subpopulation will have unique G1 and G2 peaks, and the number of subpopulations is the total number of peaks divided by two.

Conversely, if the number of peaks is odd, this indicates that the G1 peak of one subpopulation is the same as the G2 peak of another subpopulation. For example, in the case of a population that contains diploid (2N) and tetraploid (4N) subpopulations, the diploid G2 peak is the same as the tetraploid G1. In this case the number of subpopulations is the total number of peaks plus one divided by two. *findPairs()* assumes that each peak remaining after *findTruePeaks()* is a possible G1 and will try to find their subpopulation pairing. The G2 boundary conditions in *PeakAlgorithm2()* are the same as in *PeakAlgorithm1()*. For each possible G1 peak, *findPairs()* looks at all peaks identified within the specified range; if there is more than one peak identified, it will assign the tallest of those identified peaks to be the associated G2 peak. If there is no possible G2 peak within the range, the algorithm will assume that this possible G1 peak does not have a subpopulation pairing, and it will be removed from the list of potential subpopulations.

After the subpopulations have been identified, the number of cells associated with each subpopulation is determined and the percent of cells accounted for by all identified G1 and G2 peaks is calculated (*cellPerc*, (Figure **??**)). Analogous to *popBoundary*, it the proportion of unused cells is large, the algorithm will fail. A default value of 86% was set based on our test dataset, but again this is tuneable and should be evaluated by each researcher. As with *PeakAlgorithm1()*, if a sample fails, the first step is to run *doubletCheck()* to identify and take into account doublets. If *cellPerc* remains below the threshold, the data will be passed to the next algorithm, *peakAlgorithm3()*. The process starts again at the peak detection step. The interval where a G2 peak is looked for is widened to (1.7 *× G*1, 2.3 *× G*1), and *findTruePeaks()* and *findPairs()* are applied as before. If the sample’s *cellPerc* still does not pass the threshold of 86% in *peakAlgorithm3()*, then it gets passed on to *peakAlgorithm4()*, where the smoothing level is reduced to four, and *findTruePeaks()* and *findPairs()* are applied as before. Samples that fail at the end of *peakAlgorithm4()* are flagged in the final data table for user investigation and visual inspection, as there is less confidence in the identified peaks. They are nevertheless passed on to the final function, *peakAlgorithm5()*, which has a smoothing level of three and the G1 and G2 peak interval is widened to (1.5 *× G*1, 2.5 *× G*1) in a final attempt to identify possible pairs (Figure **6**).

**FIG 5.**
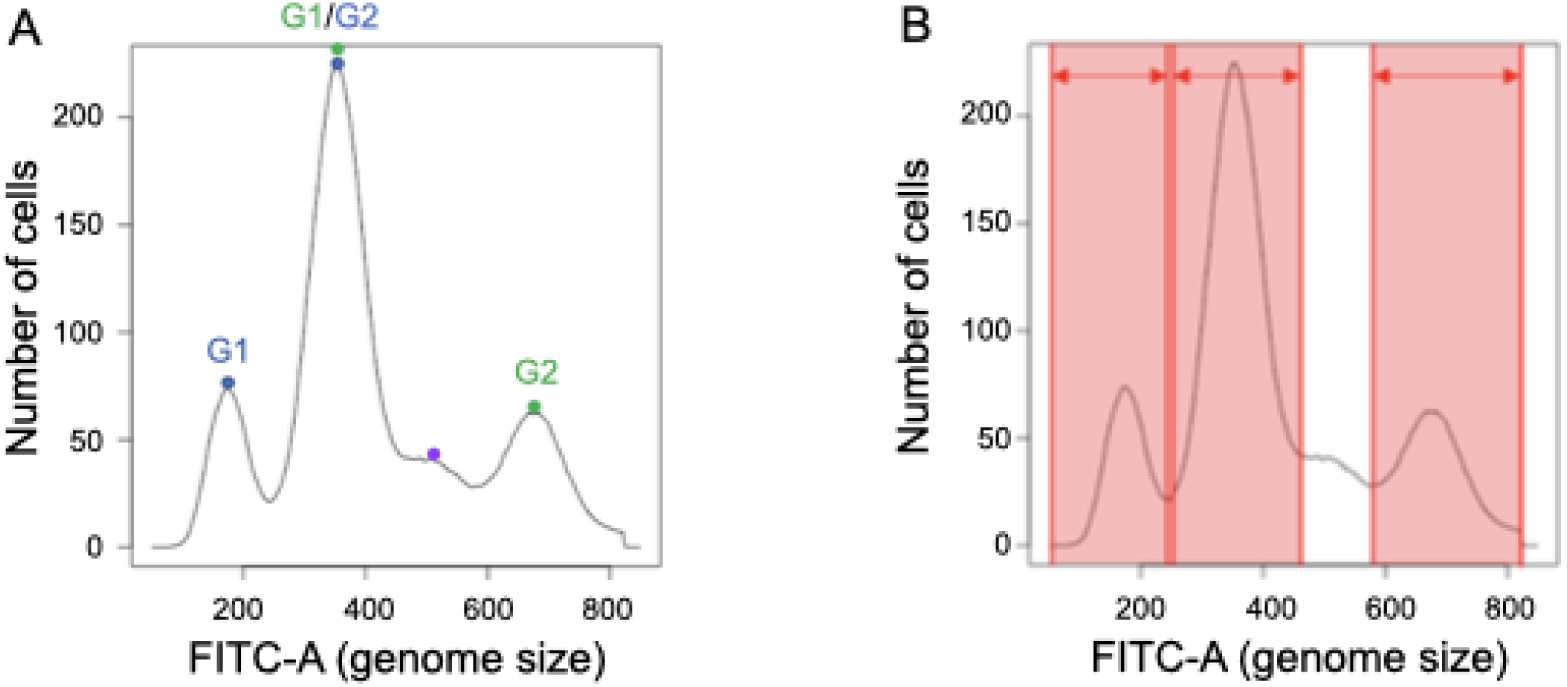
The percent of cells being used in the subpopulation pairings in a sample with two subpopulations. A) The blue dots represent the G1 and G2 peaks of the first subpopulation. The green dots represent the G1 and G2 peaks from the second subpopulation (note that this green G2 peak is not called as a doublets since it is higher than the small peak indicated by the purple dot at G1 + G2 from the blue population). B) The percent of cells being used within all subpopulation pairings is represented by the red areas, while the area in white are the cells not being used. *cellPerc* is calculated by the number of cells in the red area divided by the total number of cells *×* 100.

**FIG 6.**
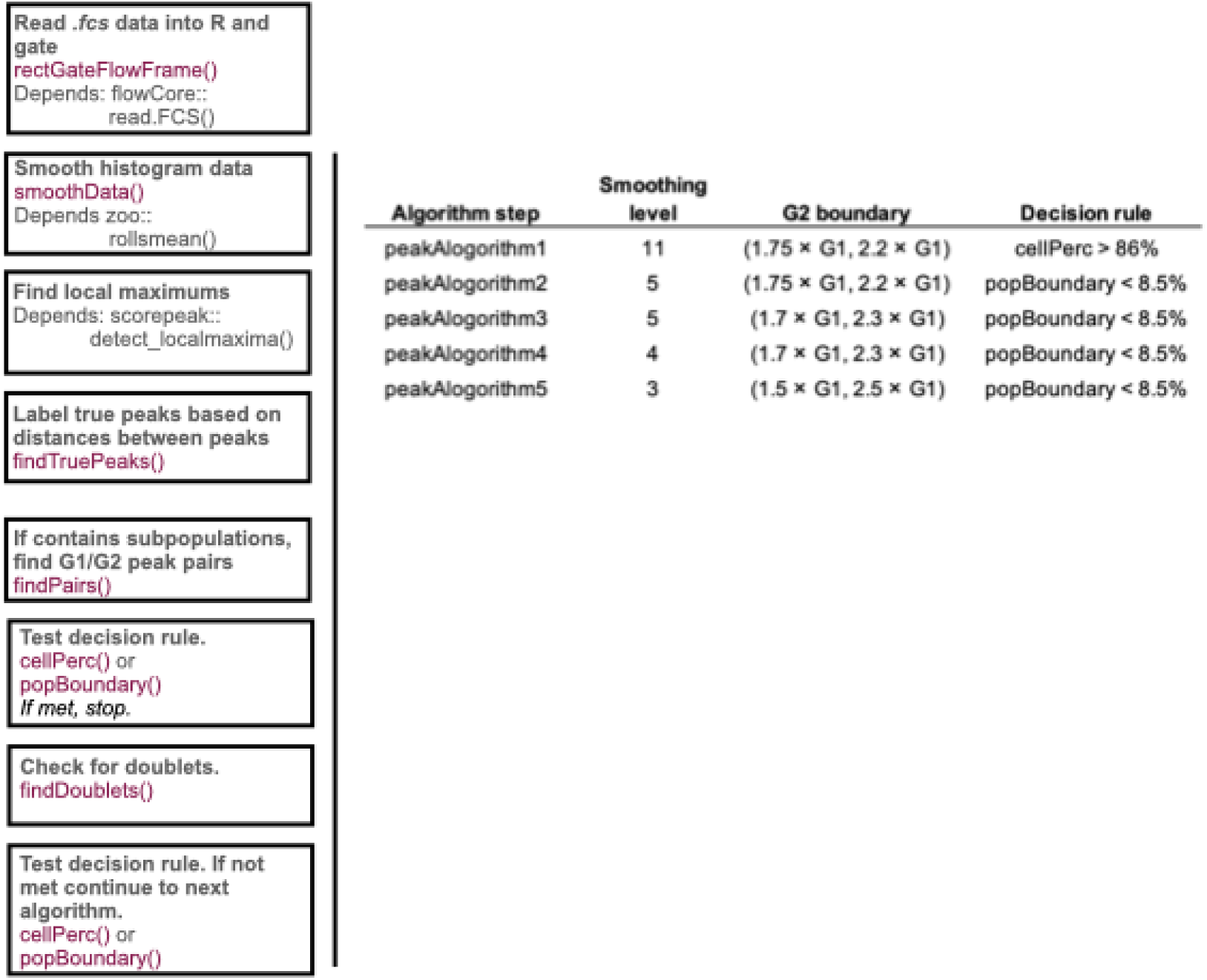
General workflow of the peak detection algorithm functions. The new functions are highlighted in purple.

### Calculating confidence with residual standard error (RSE)

Following peak detection analysis, the Dean-Jett-Fox cell cycle algorithm is applied to each sample using the identified peak means and heights (including from each subpopulation and doublet peak, where appropriate) to calculate a residual standard error (RSE) using nonlinear least squares models. Up to three DJF models are run for each sample: a single population model, a model including doublets, and a model including subpopulations.

The DJF cell cycle algorithm [25] assumes the peaks are Gaussian distributed with

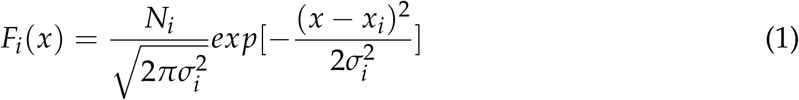

Where the *x_i_*’s are represented by the means identified in our peak algorithm. The *σ_i_*’s are calculated as the width of the distribution at 60% [24] of each mean, i.e., if the G1 mean is 170, the standard deviation will be calculated where the height is 0.6 *×* 170 = 102 on either side of the peak. The *N_i_*’s are calculated by summing the cells within (*↔σ_i_*,*σ_i_*).

The first model assumes each sample is a single population, where

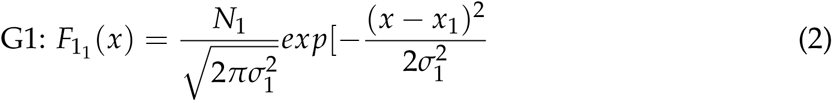

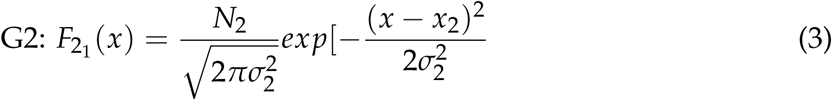

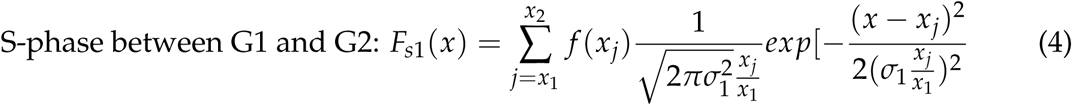

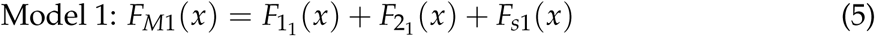

The second model adds doublet peaks to the single population, if any were identified by *doubletCheck()*.

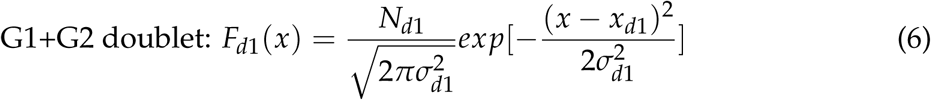

S phase between G2 and G1+G2 doublet:

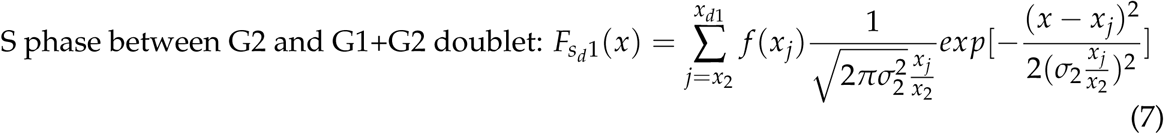

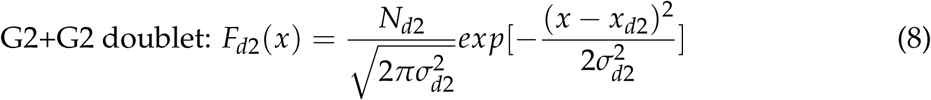

S phase between G1+G2 doublet and G2+G2 doublet:

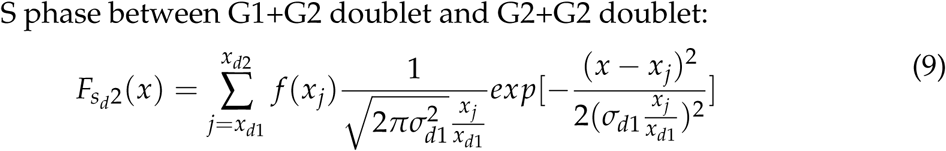

Model 2:

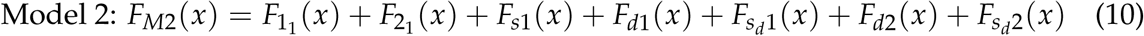

If *flowPeakDectection()* identified any subpopulations, the third model fits each subpopulation identified by the algorithm.

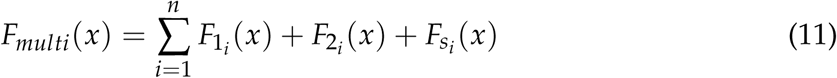

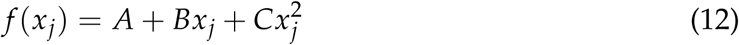

A, B and and C are estimated using nonlinear least squares.

The final RSE for the sample is determined as the lowest of the fitted models (Figure **7**).

**FIG 7.**
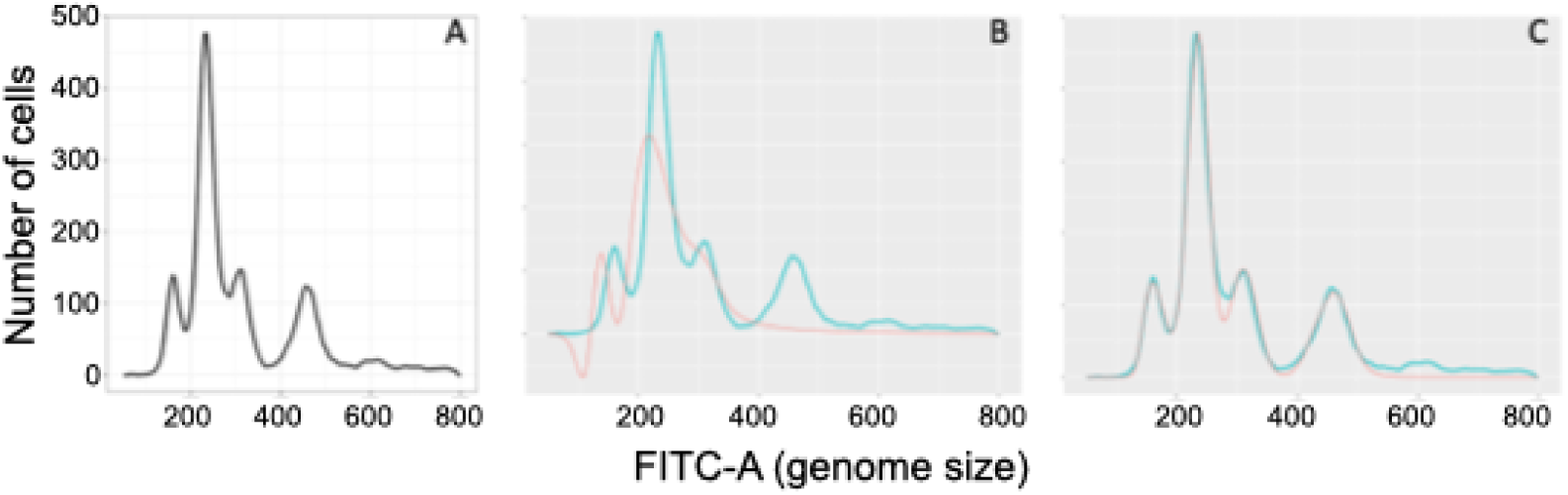
RSE Model Comparison. A) The sample we are fitting a model to, flowPeakDetection() identified two subpopulations with 4 peaks located at x=160, 235, 310, 455. B) The fit from model 1, assuming a single population. The green line is the trace of the sample of interest and the pink line the model fit, resulting in a RSE of 59.35. C) The fit from model 3 output, with an RSE of 11.85.

Users can use the RSE as a confidence term to see how well the algorithm performed relative to their visual observation or among other samples. The function *RSEOutlierDetection()* plots a histogram of the calculated provided RSE for a flow set and measures the skewness of the distribution. If the skewness is greater than zero, the function flags and plots the samples whose RSE falls in the top *ε* percent of all samples. (If the skewness is less than or equal to zero, then the function provides a message to the user and saves the histogram). The default outlier setting is to use *ε* = 0.05, but this can be adjusted based on the visualization.

### Peak correction

The final output file contains information about the location of all G1 and G2 peaks, the number of cells in each, RSE values for the three models, and the peak detection algorithm that was used. In addition, the file contains a column indicating whether doublets were identified and a column labelled ‘investigate’, which indicates to the user that a visual inspection should be conducted because either there are either an excess of cells either to the left of the first identified G1 peak, or the percentage of cells outside a boundary exceeds a specified threshold. In the event that the expected subpopulations following visual inspection does not match the ploidyPeaks model result, the user can attempt to force *ploidyPeaks* to fit a model with the desired number of subpopulations using the function *peakCorrection()*.

### Visualizations

Visual analysis is an important component of flow cytometry. *ploidyPeaks* automatically provides users with a bivariate visualization of the gating procedure for each sample. In addition, a core function *flowLineGraph()* provides a flexible method to plot single or multiple fluorescence intensity histograms together. To compare multiple samples, *flowLineGraph()* includes the option to either plot multiple samples on top of each other or to plot multiple samples into a 3 *×* 4 grid (if more than 12 samples are specified, additional pages will be added to the output). There are many flexible options, including line colour, options for vertical line placement (e.g., to show G1 means), and to add annotations (e.g. to add RSE values).

### Package documentation

An extensive vignette including package walkthrough are available at. The major package functions are listed in Table (**1**).

**TABLE 1.** Major ploidyPeak functions.

| Function | Inputs | Actions | Outputs |
| --- | --- | --- | --- |
| rectGateFlowFrame | Sample directory, sample name, $x$ and $y$ variables, numerical bounds for gating, option to save plot. | Reads in a single flow sample and applies gate. | Single *.fcs* file of gated data and plots of the data pre- and post-gating. |
| rectGateFlowSet | Sample directory, $x$ and $y$ variables, numerical bounds for gating, option to save plot(s). | Reads in a set of flow samples and applies gates to each. | Gated data *.fcs* files, plots of data pre- and post-gating, and *.csv* file containing the percent of cells gated out of each sample. |
| flowLineGraph | Sample directory, sample name(s), $x$ variable, (optional) control sample name, (optional) colour for each sample, option to save plot(s). | Plots histogram(s) on a single graph or in a grid. | Histograms are saved. |
| flowPeakDetection | Sample directory, sample name(s), $x$ variable, population thresholds, doublet max height, option to save doublet information, option to save the NLS | Uses peak detection algorithms to find local maximums and matching G1 and G2 peaks. Flags a sample if the thresholds are exceeded. | Saves a *.csv* file with peak location(s), number of cells in each peak, whether the sample was flagged, RSE and NLS fit graphs for each sample. |

### Testing dataset

Replicate populations of *C. albicans* from twenty ancestral strain backgrounds were evolved to the azole drug fluconazole (FLC) for 100 generations through experimental evolution. Twelve replicate lines from each of the ancestral strain backgrounds were passaged through 1:1000 transfers every 72 h in YPD + 1 *µ*M FLC, for a total of 240 replicates (listed as A1-A20, full strain information is in [31]). Each replicate was frozen at the beginning of the experiment (t0), after 30 generations of evolution (t3), after 80 generations (t8), and after the final timepoint (t10). Samples were prepared for flow cytometry following a standard procedure as previously described [31]; briefly, replicates were inoculated from frozen into YPD and grown overnight prior to ethanol fixation, RNAse A treatment, and overnight staining at room temperature with SYBR green. All isolates from a given time point were prepared and run on a BD Biosciences LSRII at the same time. In total, data from 960 flow cytometry samples were collected in *.fcs* files.

An analysis of the t0 and t10 replicates was previously published [31] following a manual analysis by A.C. Gerstein of the data in FlowJo. The two additional timepoints (t3 and t8) had also been manually analyzed at the same time by A.C. Gerstein, but not included in the original manuscript. All t0 samples should be euploid diploids, as the ancestral strains are all known to be diploid, and the t0 replicates were isolated from single colonies streaked onto YPD plates, where the karyotypic mutation rate should be negligible. However, once the replicates are evolved in the drug, we expected variants to arise rapidly, and potentially in some cases sweep through the population. An example of manual gating for analysis of subpopulations is shown in Figure S1. The same *.fcs* files were analyzed here with *ploidyPeaks*.

Scoring for mismatches was done by graphically comparing G1 means identified from the manual analysis (MAN) and *ploidyPeaks* (PP) results. *flowLineGraph()* was used to plot the fluorescence intensity histogram from all mismatches, using vertical lines to annotate the PP G1 calls and green dots to indicate the manual analysis (MAN) (examples in Figure 3, all mismatches in Figures S2-S4). Mismatches between the two methods were identified independently by three of the authors, with each sample scored whether PP was correct, MAN was correct, or it was unknown (UNK, erring towards UNK when it did not seem obvious). Most samples were scored consistently by all three authors, or a consensus was easily reached where there was initially a disagreement (with bias towards UNK where necessary).

When the manual analysis was correct, and the number of populations identified by PP was different, we used *peakCorrection* to try to force *ploidyPeaks* to recover the same result. We also took note of whether *ploidyPeaks* had flagged the mismatched samples, as these are samples where the program user is highly encouraged to conduct a visual inspection following computational analysis. For all samples that contained multiple populations, we determined the “major” population as the one with the most cells, with other subpopulations annotated as “minor”. For MAN samples, we based this distinction on G1 peak height, following [22]. The final dataset used to plot the timeseries data included corrected values from the manual analysis; we retained the PP values for UNK samples. There is commonly slight variation in flow cytometry from data to day even when using the exact same machine with the same settings, as we did in the collection of this dataset. To correct this machine variation among timepoints, we applied a correction factor to t3, t8, and t10 data. For each timepoint, we determined the mean diploid G1 peak intensity from all replicates evolved from A12 and A18 backgrounds, which never exhibited obvious genome size changes. We then multiplied the G1 mean at t0 divided by the G1 mean at t3, t8, or t10 for all data. The base genome size (fluorescence intensity) for diploid for visualization purposes was determined as the mean fluorescence intensity of all replicates to t0.

## RESULTS and DISCUSSION

We developed a new computational method to analyze genome size flow cytometry data. The *ploidyPeaks* R package builds off the Dean-Jett-Fox mathematical framework included in the popular program FlowJo to identify G1 and G2 means from cell cycle analysis. We extended the existing method to account for cell populations that contain not just a single genome size, but populations with up to three genome size subpopulations. In addition to fitting G1 and G2 means for each subpopulation, *ploidyPeaks* additionally calculates the residual standard error (RSE) for each sample from the single population model, a model that includes doublets, and where relevant the model that includes the identified subpopulations.

We compared how *ploidyPeaks* performed to a manual analysis of flow cytometry data conducted in FlowJo by an experienced flow cytometry user (ACG). 960 samples were previously collected and analyzed in FlowJo from replicate populations evolved from 20 ancestral strains. Twelve replicates from each strain were evolved for 100 generations in YPD + fluconazole, a drug that has previously been shown to cause genome size variation in evolved replicates. Samples were taken at four time points, t0, t3 (30 generations), t8 (80 generations), and t10 (100 generations) (Figure **8**). Out of 960 samples, 95 mismatches were identified between the Flow Jo (i.e., manual, MAN) analysis and the automated *ploidyPeaks* (PP) analysis. Over a third of them (39.0%) were flagged by *ploidyPeaks* for visual inspection. This is a much higher proportion than across the entire dataset, where only 146 samples (15%) were flagged. The distributions of mismatches among timepoints were relatively similar (excepting t0, with only five mismatched samples); there were 30, 25, and 36 mismatches for t3, t8, and t10, respectively (Table **2**). The distribution of mismatches among strain backgrounds was also variable. The five strains with an ancestral *MIC*_50_ above the evolutionary drug environment generally exhibited few mismatches, as most isolates remained diploid (as we previously showed, [31]). Many isolates (16/75) had multiple time points with mismatches, consistent with mismatches being most common for isolates with subpopulations, and some isolates having acquired genome size subpopulations early in the evolutionary trajectory and retaining them for multiple time points.

**FIG 8.**
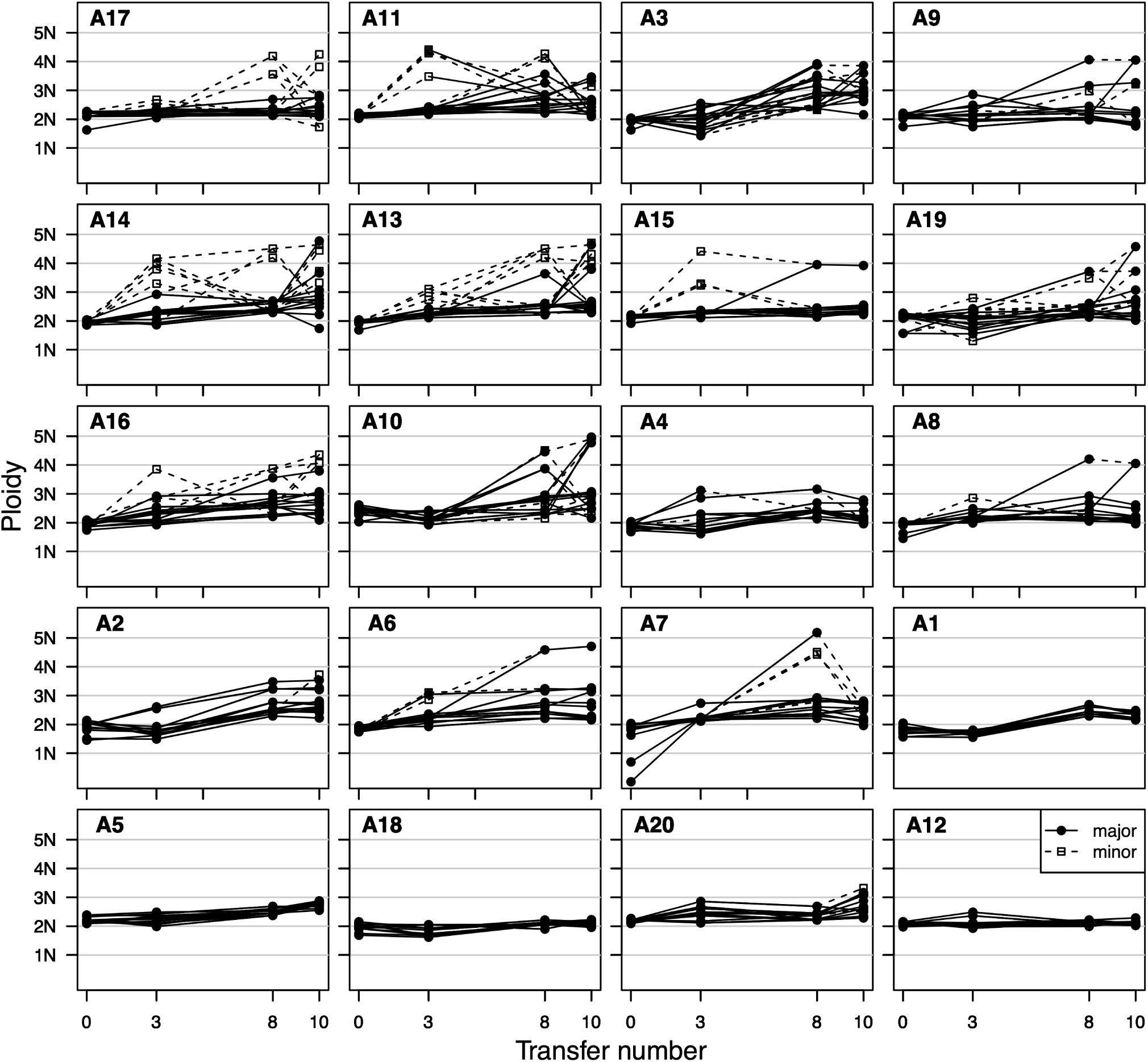
Change in genome size over time. Strains are ordered by increasing ancestral growth ability in the evolutionary environment from [31]. The labelleling of major and minor points was decided by cell counts for PP and UNK populations and a visual comparison of peak height from MAN populations.

**TABLE 2.** Mismatches between manual analysis and *ploidyPeaks*. Ancestral *MIC*_50_ was measured as previously described [31].

| Strain | t0 | t3 | t8 | t10 | total | Ancestral<br>$MIC_{50}$ |
| --- | --- | --- | --- | --- | --- | --- |
| A1 | 0 | 0 | 0 | 0 | 0 | > 1 |
| A5 | 1 | 0 | 0 | 0 | 1 | > 1 |
| A12 | 0 | 0 | 0 | 0 | 0 | > 1 |
| A18 | 0 | 3 | 0 | 0 | 3 | > 1 |
| A20 | 0 | 0 | 0 | 5 | 5 | > 1 |
| A2 | 0 | 1 | 0 | 2 | 3 | = 1 |
| A6 | 0 | 2 | 0 | 1 | 3 | = 1 |
| A7 | 2 | 0 | 2 | 2 | 6 | = 1 |
| A3 | 0 | 1 | 4 | 1 | 6 | < 1 |
| A4 | 0 | 1 | 0 | 1 | 2 | < 1 |
| A8 | 0 | 0 | 1 | 1 | 2 | < 1 |
| A9 | 0 | 0 | 2 | 3 | 5 | < 1 |
| A10 | 0 | 1 | 4 | 2 | 7 | < 1 |
| A11 | 0 | 4 | 2 | 1 | 7 | < 1 |
| A13 | 1 | 2 | 1 | 3 | 7 | < 1 |
| A14 | 0 | 4 | 6 | 5 | 15 | < 1 |
| A15 | 0 | 4 | 0 | 0 | 4 | < 1 |
| A16 | 0 | 3 | 2 | 3 | 8 | < 1 |
| A17 | 0 | 2 | 0 | 5 | 7 | < 1 |
| A19 | 0 | 2 | 1 | 1 | 4 | < 1 |
| **Total** | **4** | **30** | **25** | **36** | **95** |  |

Having identified 95 mismatched samples, we next sought to determine whether the MAN or PP analysis was correct. All mismatched samples were plotted using *flowLineGraph()* and visually inspected. Each researcher assigned a visual class depending on which analysis was deemed to have done a better job of peak identification: PP, MAN or unknown/inconclusive (UNK). We classified that PP was likely correct for 35 samples, the MAN analysis was likely correct for 23 samples, and 37 samples were unknown (Figure **9**A.) Representative traces with called peaks from each class are provided in Figure **9**B-D. We were encouraged to see only nine samples where the manual analysis was assessed as correct that were not flagged (an error rate well below 1% across the entire dataset). Accross all samples, the manual analysis had identified an additional subpopulation compared to *ploidyPeaks* in all but two cases (Figure S2). When we ran *peakCorrection()* on these subsamples (forcing an additional peak), it successfully identified the same peaks as the manual analysis in half of the cases (12/21) (Table S1).

**FIG 9.**
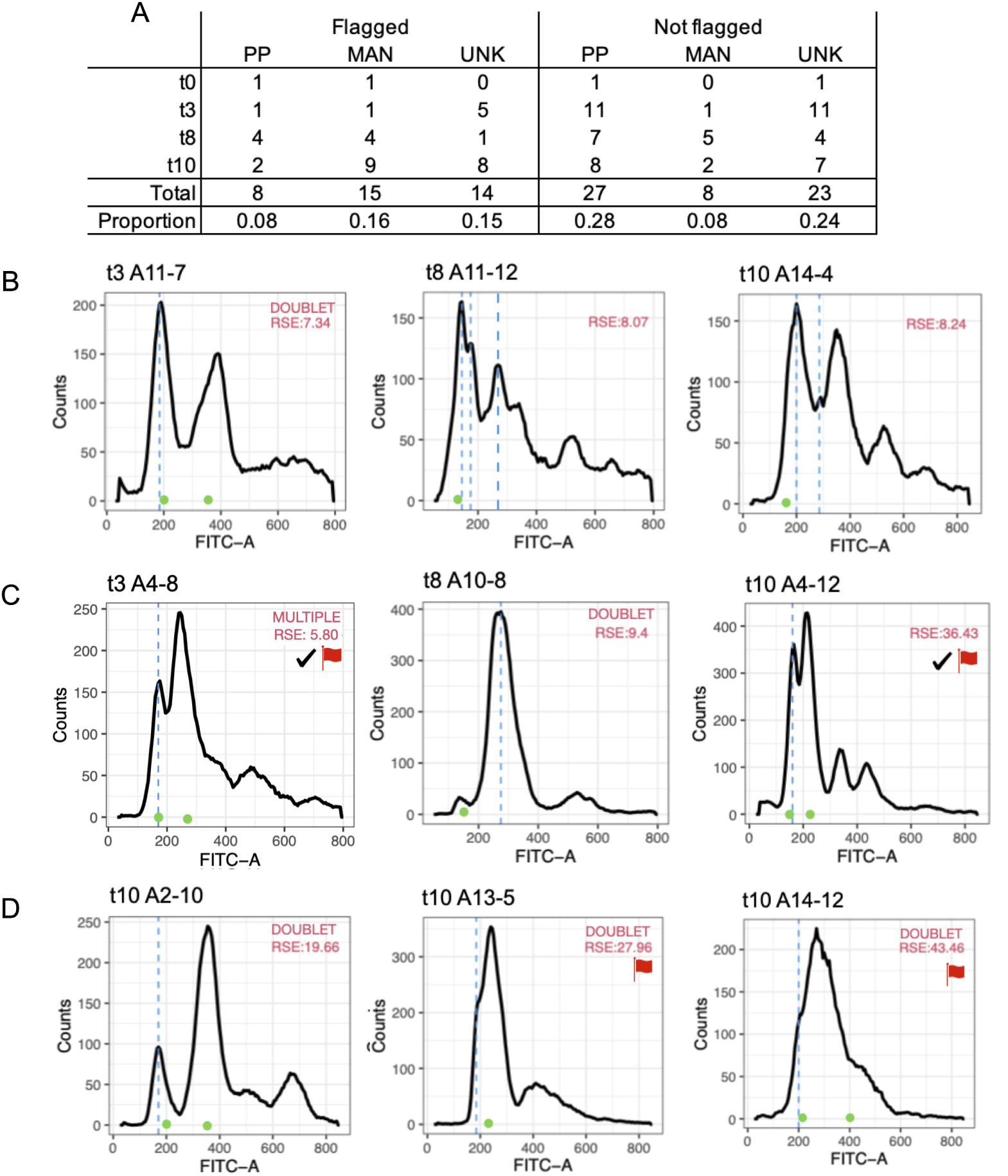
Mismatch analysis. A) Overview of mismatch results, split by whether the sample was flagged by *ploidyPeaks*, PP: *ploidyPeaks* deemed correct, MAN: manual analysis deemed correct, UNK: unknown. Example samples where we deemed B) PP correct, C) MAN correct or D) UNK. Red flags indicate whether *ploidyPeaks* indicated the sample should be examined, a checkmark indicate samples where secondary analysis with *peakCorrection* identified the same peaks as the manual peaks, arrows on the x-axis indicate the peaks identified by *peakCorrection* when it did not match.

The 35 samples where we thought *ploidyPeaks* was correct were flagged much less often than the other cases. Of the eight samples that were flagged, five had multiple populations identified by *ploidyPeaks* but not the manual analysis, while the other two had a single peak placed in different locations. The non-flagged samples were similarly distributed, near equal split among samples with extra peaks identified in either the manual or *ploidyPeaks* analysis, with a minority of samples having a single G1 mean in axdifferent place (Figure S3). This speaks to the original motivation in the development of *ploidyPeaks*. In order to identify multiple G1 peaks with a manual analysis, gates have to be individually drawn around the predicted subsamples through intuition.

Looking at the inconclusive samples, a few patterns emerge. Fifteen of the inconclusive samples are from t10, while 16 are from t3 and only five from t8. Although it is possible that something about the period of evolution in fluconazole influences the likelihood that a sample will be difficult to analyze, it also seems likely that the timepoint variation is driven by something that is not biological, but rather something related to either the sample prep or flow cytometry machine variation (e.g., the type of samples that were previously run on the machine). Some of the samples, particularly from certain strain backgrounds (A3, A14), did not exhibit the characteristic double-normal distribution peak required to fit cell cycle models correctly (Figure S4). Compared to the manual analysis, in order to call a G1 peak *ploidyPeaks* has a much stricter requirement for there to be an identified and appropriate peak (i.e., local maxima) in the correct G2 range. There are also a number of cases where it did not seem possible to assess whether high genome size peaks were simply doublets (causing a peak at G2+G2), or an endoduplicated subpopulation (with a G1 peak at the initial population G2 peak). The default setting for the maximum height for doublets in *ploidyPeaks* was based on test data generated from mixtures of known euploids, and experience knowing the characteristic pattern of peak heights, i.e., that the G2 peak is rarely larger than the G1 peak. This knowledge led us to decide on how to call the presence of doublets, based on the height of peaks at G1+G2 and G2+G2 peaks. However, we have also observed cases where the single G2 peak does seem to be significantly higher than the G1 peak, which could be biologically meaningful if cells are getting delayed through G2/M, and would have implications on the ratio of G1+G2/G2+G2 heights. We chose to implement a blanket parameter for the maximum height allowed for the G2+G2 to be called as doublets, but in the future, the analysis could be done on a sample-by-sample basis. Fourteen of the inconclusive samples were flagged by *ploidyPeaks* (Figure S4). Ideally, users will make a direct comparison of the peaks called alongside visual inspection of the genome size variation histograms to ensure that *ploidyPeaks* is consistent with their visualization and intuition for all samples.

Having established a method to identify subpopulations with less bias than the manual approach, we believe we can now more confidently ask biological questions about the stability of karyotypic variation. A visualiation of the timeseries data shows that subpopulations of higher genome size arise relatively frequently, yet only rarely are they stable or take over the population (Figure **8**). This with an analysis of a well-cited study in this space from a *Saccharomcyes cerevisiae* exeprimental evolution study that evolved replicates to heat stress [32]. The original paper described that a chromosome III aneuploidy arose and swept through the population by 450 generations, yet was replaced by genomic mutations after 1500 generations. However, a later re-analysis found that the eventual euploid populations that remained at the end were distinct from the aneuploid lineages, with aneuploidy acting as a diversion rather than stepping stone during the evolutionary trajectory [33].

We have developed an approach to analyze DNA fluorescence intensity data in an unbiased manner, through open source software that requires only minimal coding. The goals in the development of *ploidyPeaks* were to eliminate the need to rely on proprietary software, to reduce bias in the manual analysis of flow cytometry data, and to automate analysis of large datasets. In creating this package, we hope this spurs additional research into the long term dynamics and evolutionary consequenes of karyotypic variation.

## ACKNOWLEDGMENTS

We thank the many beta-testers from the Gerstein Microstats Lab who assisted in the development of ploidyPeaks package.

## DATA AVAILABILITY STATEMENT

The *ploidyPeaks* R package is available to download from GitHub at https://github.com/MicroStatsLab/PloidyPeaks.

## SUPPLEMENTAL MATERIAL

### Figures

**Figure S1. Manual gating workflow in flowJo**. A) Scatterplot of fluroescence by forward scatter, B) histogram of fluroescence of the entire population. C) Example manual gating of two subpopulations (gates drawn in pink and brown). D) Resuslting scatterplot and histogram from the two supopulations.

**Figure S2. The 23 isolates where a visual inspection concluded that the manual analysis was more likely to be correct**. The blue dashed lines indicate G1 peaks called initially by *ploidyPeaks*, green circles indicate G1 peaks called by the manual analysis. Red flags indicate the samples that were flagged by *ploidyPeaks*, and hence would have been examined following computational analysis. Checkmarks indicate samples where *peakCorrection()* identified the same peaks as the manual analysis. Arrows on the x-axis indicate the peaks identified by *peakCorrection()* when it did not match the manual analysis. The RSE shown is the lowest RSE among tested models (single or multiple peaks, with or without acconting for doublets).

**Figure S3. The 35 isolates where a visual inspection was conclude that *ploidyPeaks* was more likely to be correct.** The blue dashed lines indicate G1 peaks called by *ploidyPeaks*, green circles indicate G1 peaks called by the manual analysis. Red flags indicate the samples that were flagged by *ploidyPeaks*, and hence would have been examined following computational analysis. Checkmarks indicate samples where *peakCorrection()* identified the same peaks as the manual analysis. Arrows indicate the peaks identified by *peakCorrection()* when it did not match the manual analysis. The RSE shown is the lowest RSE among tested models (single or multiple peaks, with or without acconting for doublets).

Figure S4. The 37 isolates where a visual inspection was unable to conclude whether the manual analysis or *ploidyPeaks* was more likely to be correct. The blue dashed lines indicate G1 peaks called by *ploidyPeaks*, green circles indicate G1 peaks called by the manual analysis. Red flags indicate the samples that were flagged by *ploidyPeaks*, and hence would have been examined following computational analysis. Checkmarks indicate samples where *peakCorrection()* identified the same peaks as the manual analysis. Arrows indicate the peaks identified by *peakCorrection()* when it did not match the manual analysis. The RSE shown is the lowest RSE among tested models (single or multiple peaks, with or without acconting for doublets).

### Tables

**Table S1. Comparison of ploidyPeak output of samples where the manual analysis was correct**. Comparison of output before and after *peakCorrection()*. The column ‘Best fit model’ refers to RSE output, ‘pC’ is following *peakCorrection()*.

