## Supplemental Figures 1-4 for "PloidyPeaks: R package for automated flow cytometry cell cycle analysis and visualization in fungal microbes"

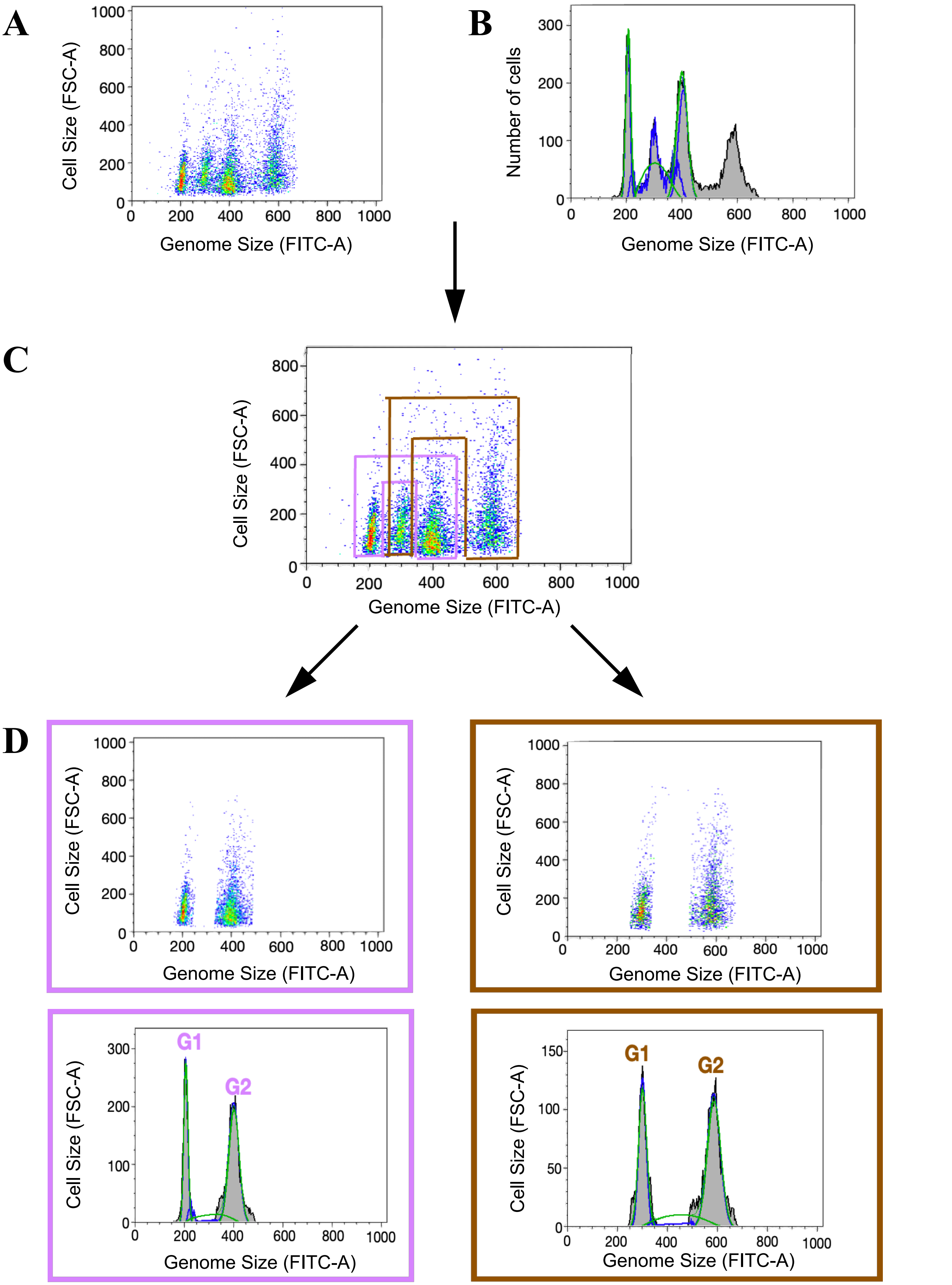


**Figure S1. Manual gating workflow in flowJo**. A) Scatterplot of fluroescence by forward scatter, B) histogram of fluroescence of the entire population. C) Example manual gating of two subpopulations (gates drawn in pink and brown). D) Resusling scatterplot and histogram from the two supopulations.

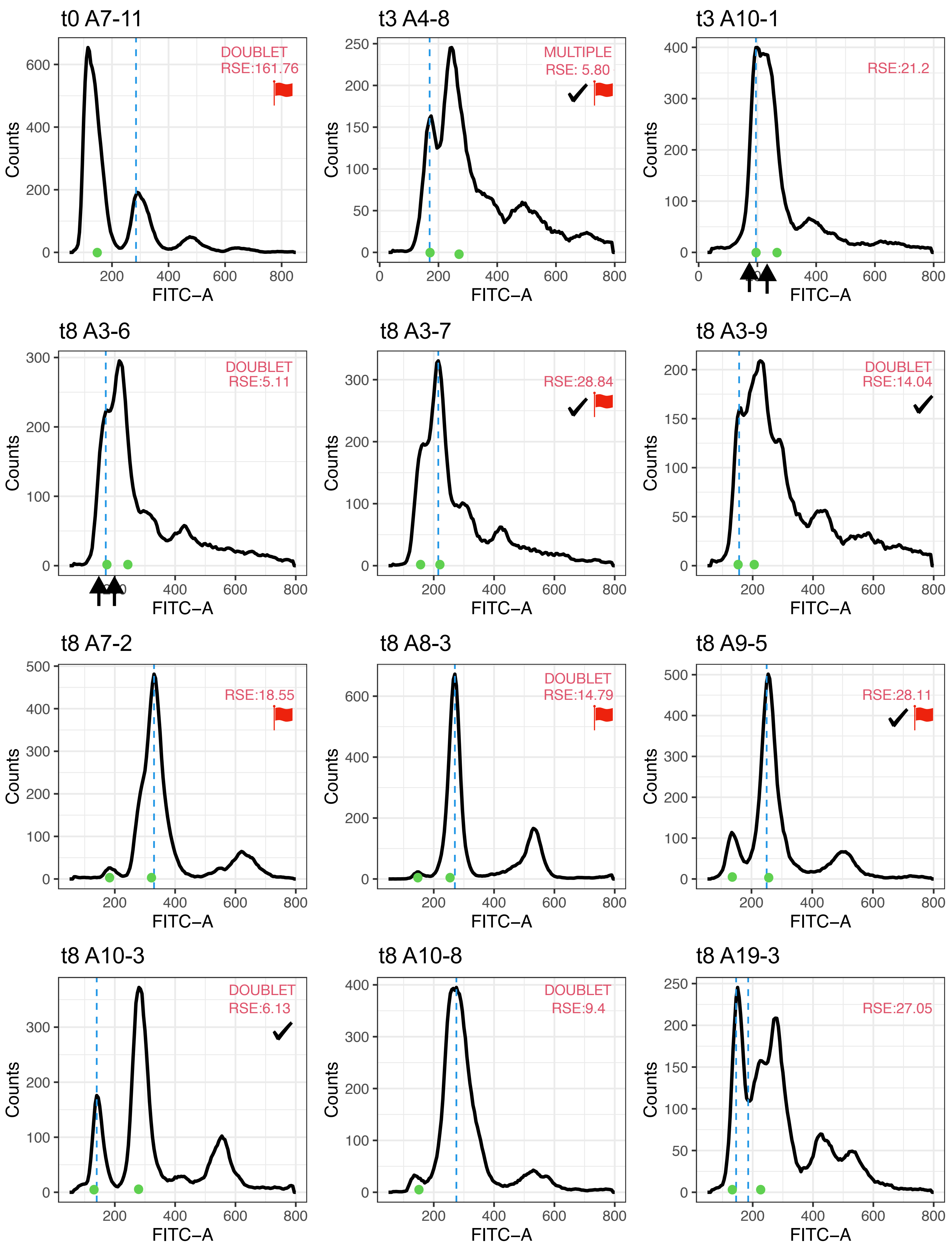

t0 A7-11
DOUBLET
RSE:161.76
t3 A4-8
MULTIPLE
RSE: 5.80
t3 A10-1
RSE:21.2
t8 A3-6
DOUBLET
RSE:5.11
t8 A3-7
RSE:28.84
t8 A3-9
DOUBLET
RSE:14.04
t8 A7-2
RSE:18.55
t8 A8-3
DOUBLET
RSE:14.79
t8 A9-5
RSE:28.11
t8 A10-3
DOUBLET
RSE:6.13
t8 A10-8
DOUBLET
RSE:9.4
t8 A19-3
RSE:27.05
Counts
FITC−A

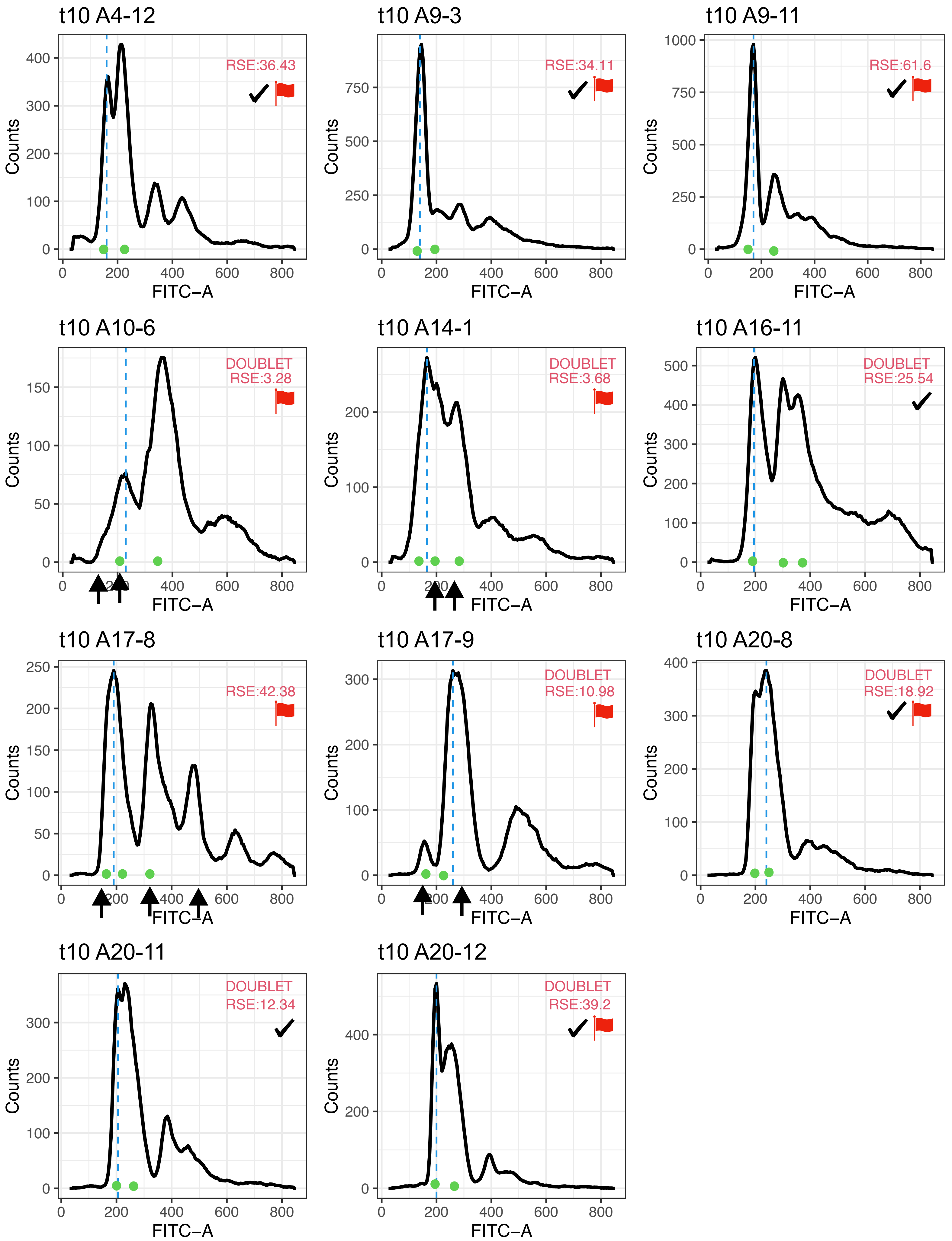
t10 A4-12
RSE:36.43
t10 A9-3
RSE:34.11
t10 A9-11
RSE:61.6
t10 A10-6
DOUBLET
RSE:3.28
t10 A14-1
DOUBLET
RSE:3.68
t10 A16-11
DOUBLET
RSE:25.54
t10 A17-8
RSE:42.38
t10 A17-9
DOUBLET
RSE:10.98
t10 A20-8
DOUBLET
RSE:18.92
t10 A20-11
DOUBLET
RSE:12.34
t10 A20-12
DOUBLET
RSE:39.2
Counts
FITC−A

**Figure S2**. The 23 isolates where a visual inspection concluded that the manual analysis was more likely to be correct. The blue dashed lines indicate G1 peaks called initially by *ploidyPeaks*, green circles indicate G1 peaks called by the manual analysis. Red flags indicate the samples that were flagged by *ploidyPeaks,* and hence would have been examined following computational analysis. Checkmarks indicate samples where *peakCorrection()* identified the same peaks as the manual analysis. Arrows on the x-axis indicate the peaks identified by *peakCorrection()* when it did not match the manual analysis. The RSE shown is the lowest RSE among tested models (single or multiple peaks, with or without acconting for doublets).

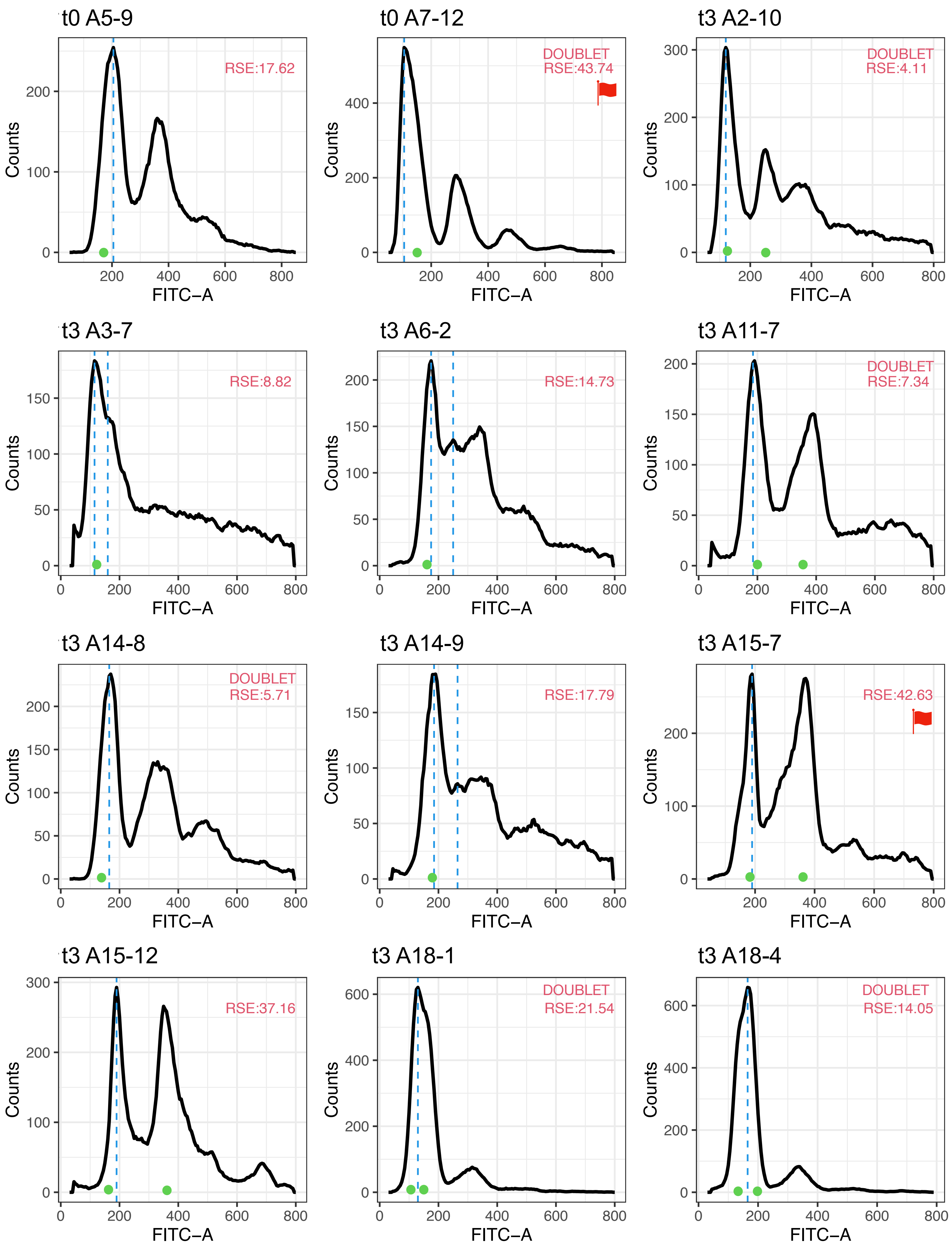
t0 A5-9
RSE:17.62
t0 A7-12
DOUBLET
RSE:43.74
t3 A2-10
DOUBLET
RSE:4.11
t3 A3-7
RSE:8.82
t3 A6-2
RSE:14.73
t3 A11-7
DOUBLET
RSE:7.34
t3 A14-8
DOUBLET
RSE:5.71
t3 A14-9
RSE:17.79
t3 A15-7
RSE:42.63
t3 A15-12
RSE:37.16
t3 A18-1
DOUBLET
RSE:21.54
t3 A18-4
DOUBLET
RSE:14.05
Counts
FITC−A

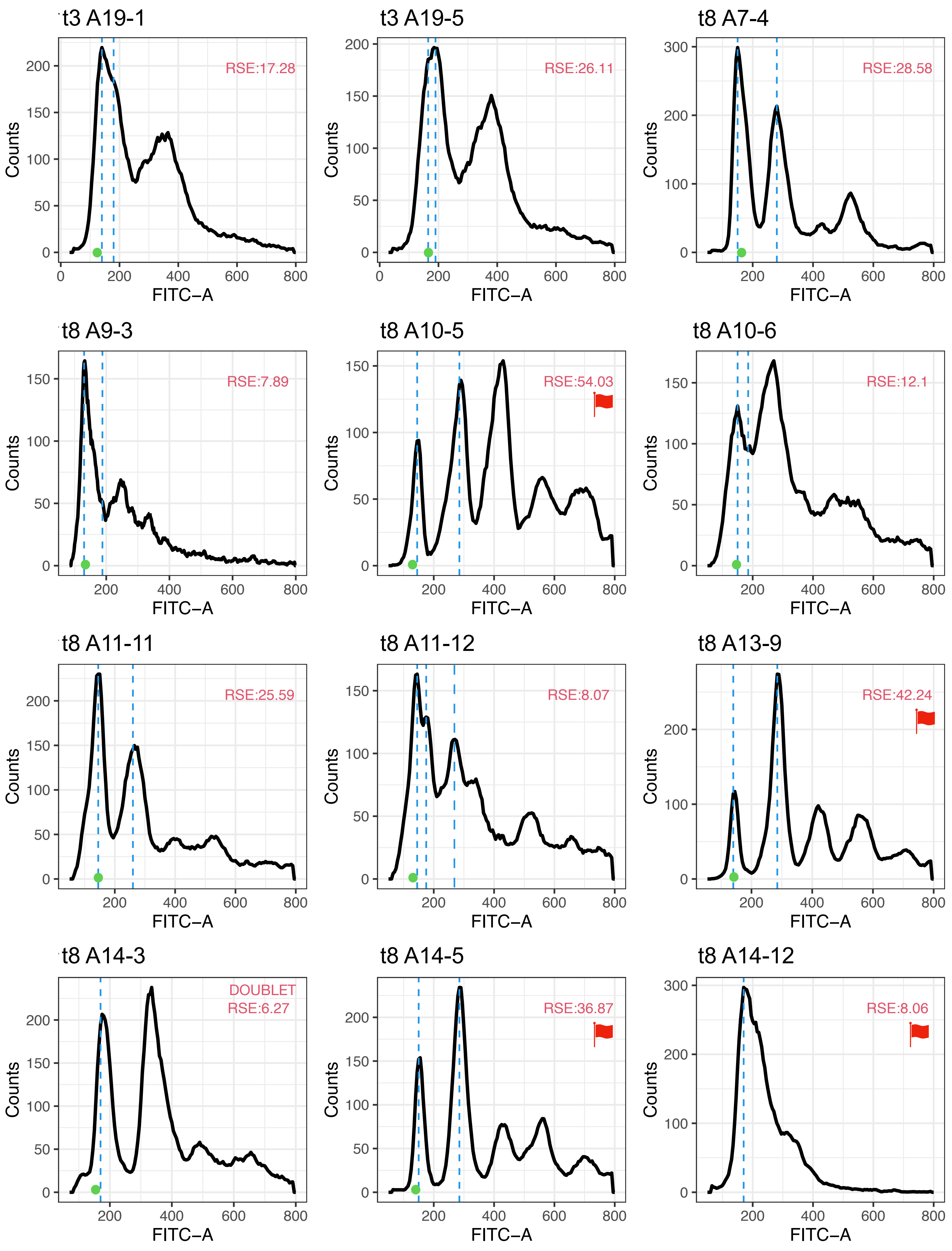

t3 A19-1
RSE:17.28
t3 A19-5
RSE:26.11
t8 A7-4
RSE:28.58
t8 A9-3
RSE:7.89
t8 A10-5
RSE:54.03
t8 A10-6
RSE:12.1
t8 A11-11
RSE:25.59
t8 A11-12
RSE:8.07
t8 A13-9
RSE:42.24
t8 A14-3
DOUBLET
RSE:6.27
t8 A14-5
RSE:36.87
t8 A14-12
RSE:8.06
Counts
FITC−A

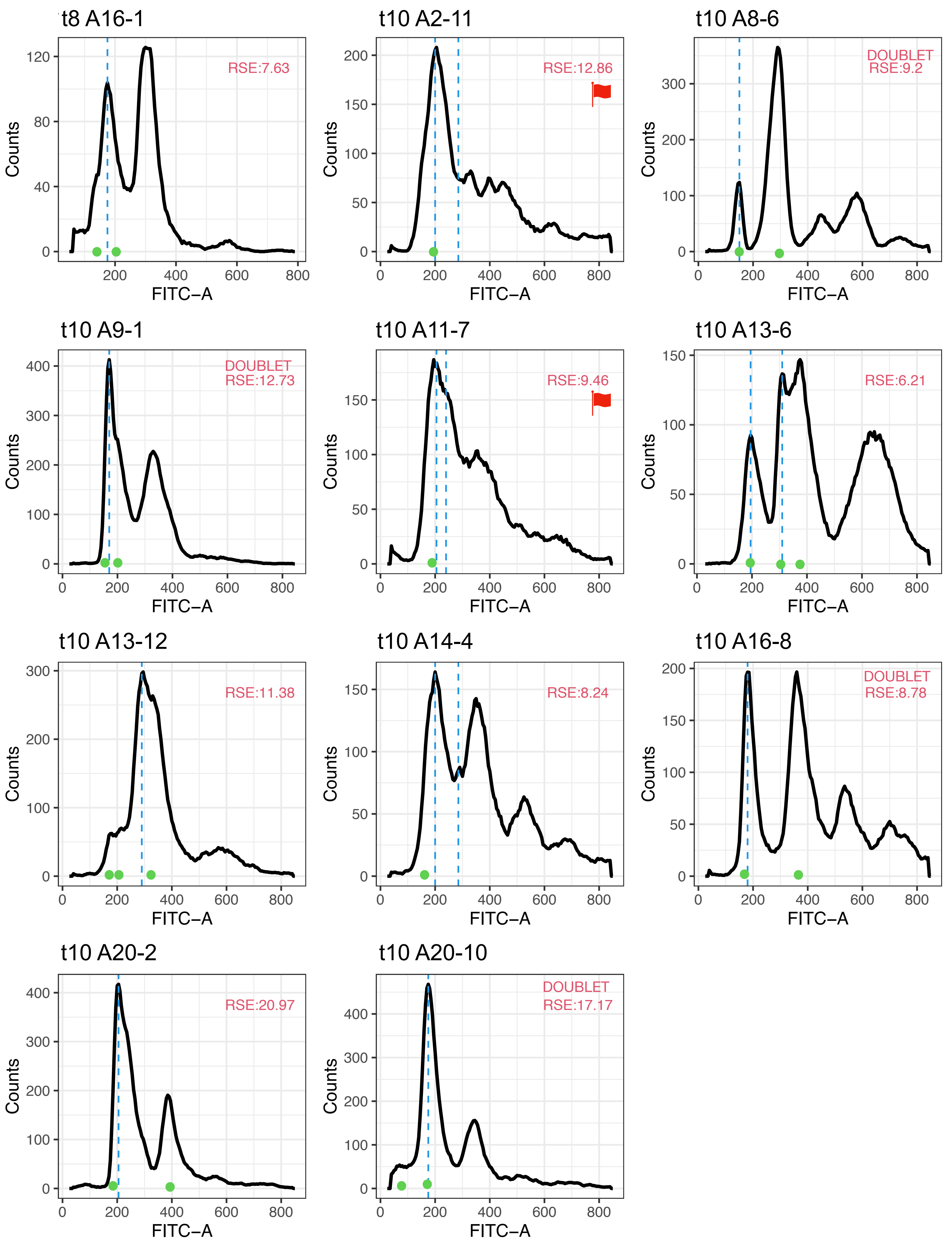
t8 A16-1
RSE:7.63
t10 A2-11
RSE:12.86
t10 A8-6
DOUBLET
RSE:9.2
t10 A9-1
DOUBLET
RSE:12.73
t10 A11-7
RSE:9.46
t10 A13-6
RSE:6.21
t10 A13-12
RSE:11.38
t10 A14-4
RSE:8.24
t10 A16-8
DOUBLET
RSE:8.78
t10 A20-2
RSE:20.97
t10 A20-10
DOUBLET
RSE:17.17
Counts
FITC−A

**Figure S3**. The 35 isolates where a visual inspection was conclude that *ploidyPeaks* was more likely to be correct. The blue dashed lines indicate G1 peaks called by *ploidyPeaks*, green circles indicate G1 peaks called by the manual analysis. Red flags indicate the samples that were flagged by *ploidyPeaks,* and hence would have been examined following computational analysis. Checkmarks indicate samples where *peakCorrection()* identified the same peaks as the manual analysis. Arrows indicate the peaks identified by *peakCorrection()* when it did not match the manual analysis. The RSE shown is the lowest RSE among tested models (single or multiple peaks, with or without acconting for doublets).

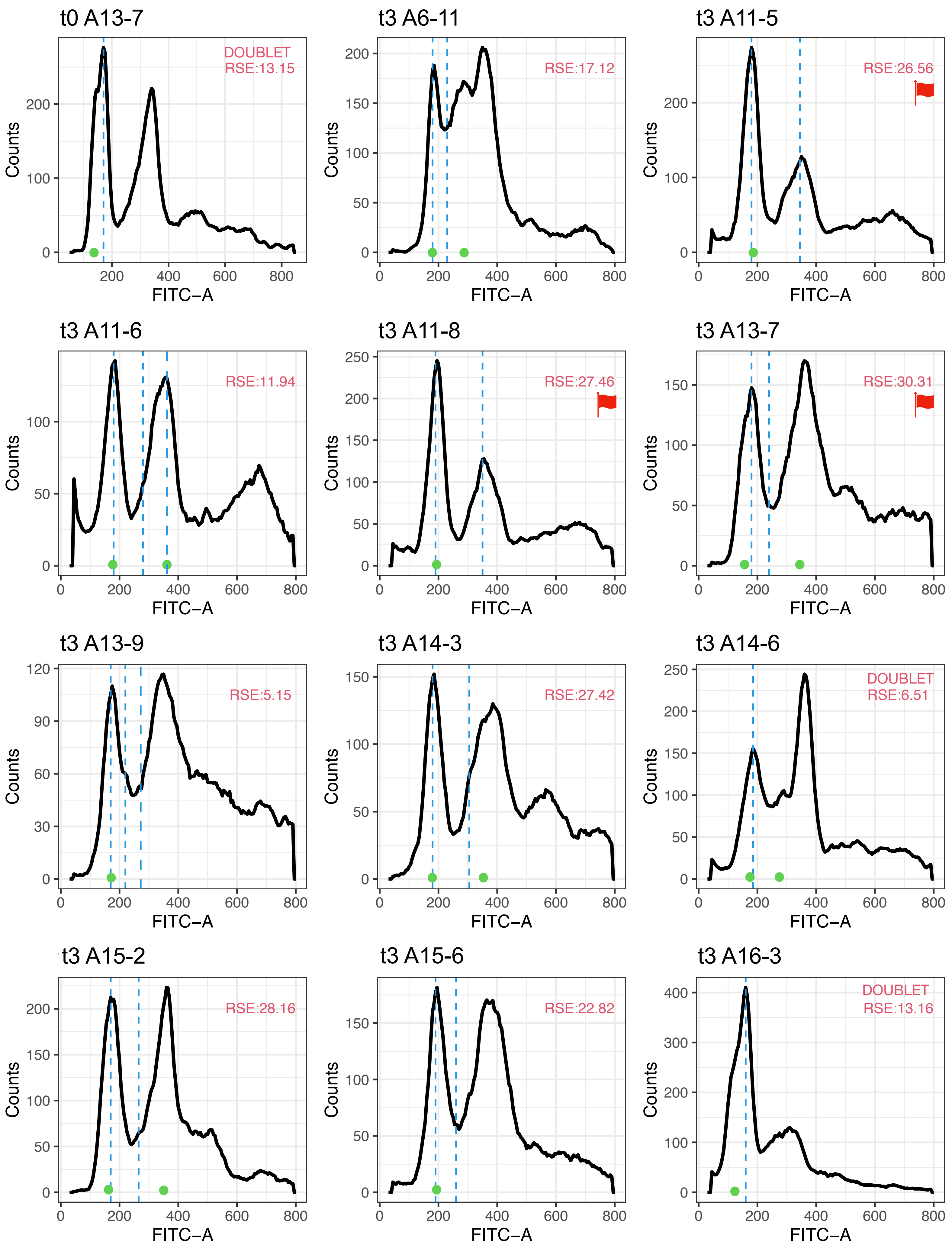
t0 A13-7
DOUBLET
RSE:13.15
t3 A6-11
RSE:17.12
t3 A11-5
RSE:26.56
t3 A11-6
RSE:11.94
t3 A11-8
RSE:27.46
t3 A13-7
RSE:30.31
t3 A13-9
RSE:5.15
t3 A14-3
RSE:27.42
t3 A14-6
DOUBLET
RSE:6.51
t3 A15-2
RSE:28.16
t3 A15-6
RSE:22.82
t3 A16-3
DOUBLET
RSE:13.16
Counts
FITC−A

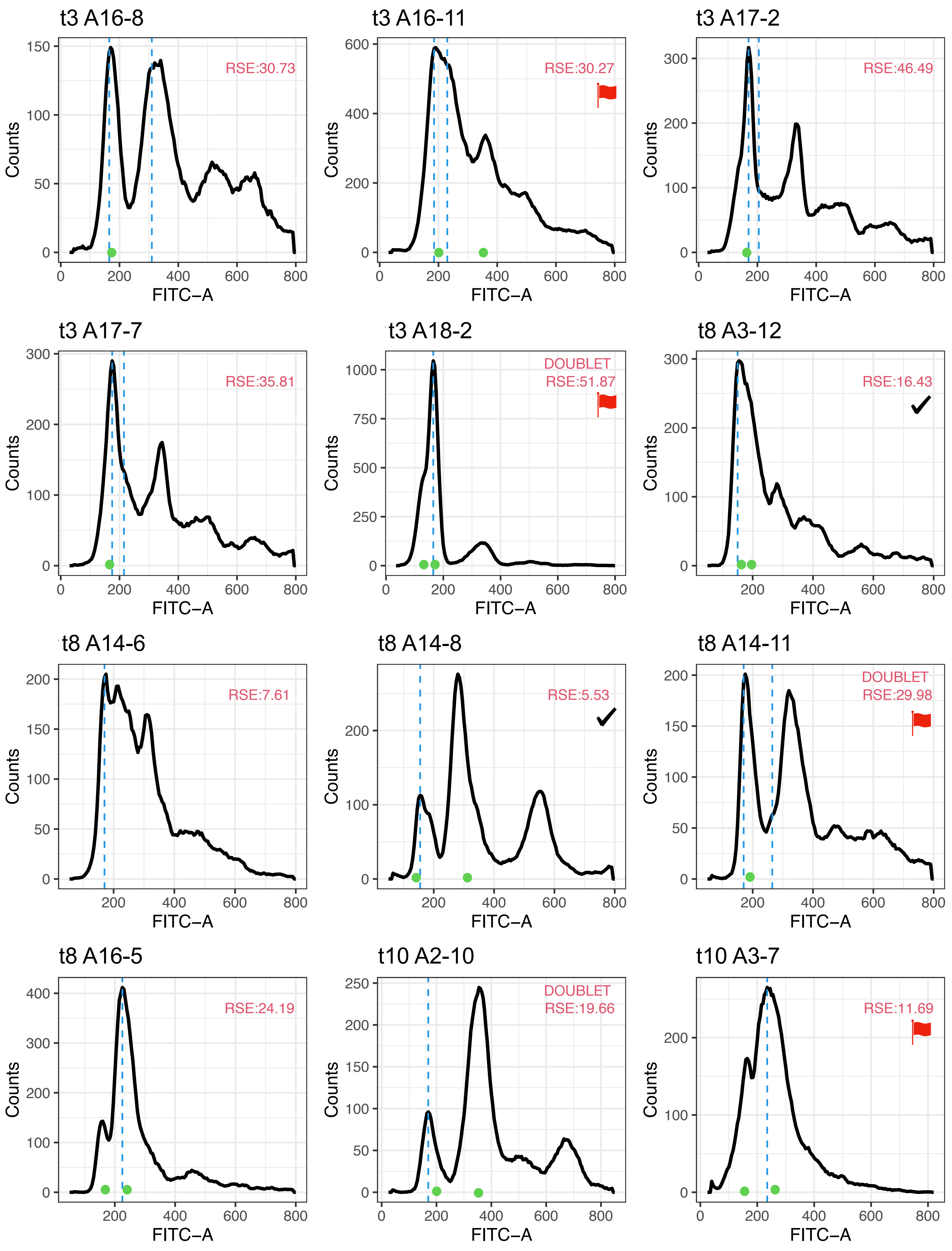
t3 A16-8
RSE:30.73
t3 A16-11
RSE:30.27
t3 A17-2
RSE:46.49
t3 A17-7
RSE:35.81
t3 A18-2
DOUBLET
RSE:51.87
t8 A3-12
RSE:16.43
t8 A14-6
RSE:7.61
t8 A14-8
RSE:5.53
t8 A14-11
DOUBLET
RSE:29.98
t8 A16-5
RSE:24.19
t10 A2-10
DOUBLET
RSE:19.66
t10 A3-7
RSE:11.69
Counts
FITC−A

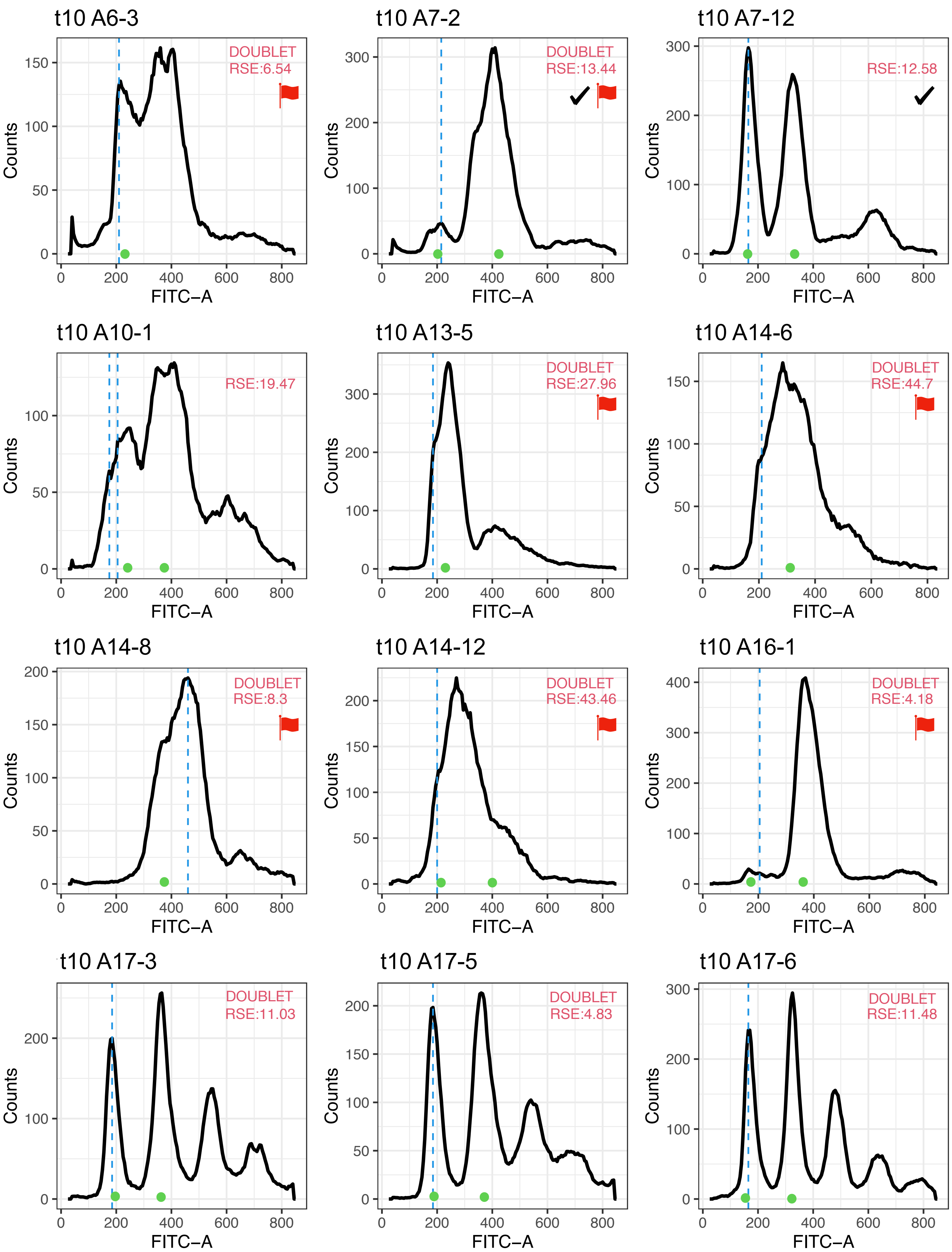

t10 A6-3
DOUBLET
RSE:6.54
t10 A7-2
DOUBLET
RSE:13.44
t10 A7-12
RSE:12.58
t10 A10-1
RSE:19.47
t10 A13-5
DOUBLET
RSE:27.96
t10 A14-6
DOUBLET
RSE:44.7
t10 A14-8
DOUBLET
RSE:8.3
t10 A14-12
DOUBLET
RSE:43.46
t10 A16-1
DOUBLET
RSE:4.18
t10 A17-3
DOUBLET
RSE:11.03
t10 A17-5
DOUBLET
RSE:4.83
t10 A17-6
DOUBLET
RSE:11.48
Counts
FITC−A

t10 A19-7

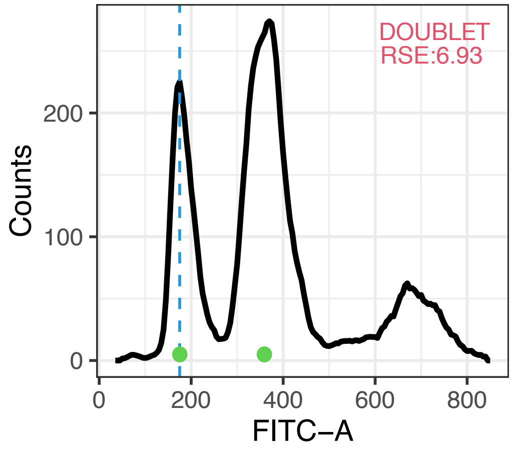


A20_11-t10

**Figure S4**. The 37 isolates where a visual inspection was unable to conclude whether the manual analysis or ploidyPeaks was more likely to be correct. The blue dashed lines indicate G1 peaks called by *ploidyPeaks*, green circles indicate G1 peaks called by the manual analysis. Red flags indicate the samples that were flagged by *ploidyPeaks,* and hence would have been examined following computational analysis. Checkmarks indicate samples where *peakCorrection()* identified the same peaks as the manual analysis. Arrows indicate the peaks identified by *peakCorrection()* when it did not match the manual analysis. The RSE shown is the lowest RSE among tested models (single or multiple peaks, with or without acconting for doublets).
